# Benchmarking informatics approaches for virus discovery: Caution is needed when combining *in silico* identification methods

**DOI:** 10.1101/2023.08.07.552334

**Authors:** Bridget Hegarty, James Riddell V, Eric Bastien, Kathryn Langenfeld, Morgan Lindback, Jaspreet S. Saini, Anthony Wing, Jessica Zhang, Melissa Duhaime

## Abstract

**Background:** The identification of viruses from environmental metagenomic samples using informatics tools has offered critical insights in microbiome studies. However, it remains difficult for researchers to know for their specific study which tool(s) and settings are best suited to maximize capture of viruses while minimizing false positives. Studies are increasingly combining multiple tool outputs attempting to recover more viruses, but no combined approach has been benchmarked for accuracy. Here, we benchmarked 63 viral identification ‘rulesets’ against mock metagenomes composed of publicly available viral, bacterial, archaeal, fungal, and protist sequences. These rulesets are based on combinations of four single-tool rules and two multi-tool tuning rules. We applied these rulesets to various aquatic metagenomes and filtering strategies to evaluate the impact of habitat and viral enrichment on individual and combined tool performance. We provide a packaged pipeline for researchers that want to replicate our process.

**Results:** We found that combining rules increased viral recall, but at the expense of increased false positives. Six of the 63 combinations tested had equivalent accuracies to the highest one (MCC=0.77, p_adj_ ≥ 0.05). All of the six high accuracy rulesets included VirSorter2, five included our “tuning removal” rule, and no high performing rulesets used more than four of our six rules. DeepVirFinder, VIBRANT, and VirSorter were each found once in these high accuracy rulesets, but never in combination with each other. Our validation suggests that the MCC plateau at 0.77 is caused by inaccurate labeling of the data that viral identification tools rely on for training and validation. In the aquatic metagenomes, our “highest MCC” ruleset identified a higher proportion of viral sequences in the virus-enriched samples (44-46%) than the non-enriched, cellular metagenomes (7-19%).

**Conclusion:** While improved algorithms may lead to more accurate viral identification tools, this should be done in tandem with curating accurately labeled viral gene and sequence databases. For most applications, we recommend the use of the ruleset that uses VirSorter2 and our empirically derived tuning removal rule. By providing a rigorous overview of the behavior of *in silico* viral identification strategies, our findings guide the use of existing viral identification tools and offer a blueprint for feature engineering of new tools that will lead to higher-confidence viral discovery in microbiome studies.

## Background

Viruses are an essential component of microbial ecosystems: they influence nutrient cycling and microbial community dynamics^1^, account for 20-40% of microbial mortality per day^2^, reprogram their hosts’ metabolism,^3,4^ and horizontally transfer genes between host populations.^5,6^ The primary approach used to discover and describe viral diversity is culture-independent metagenomic sequencing. However, viral sequences remain challenging to differentiate from non-viral ones because viruses have no universal marker gene,^7^ high mutation rates,^8,9^ and relatively small reference databases relative to the magnitude of their diversity.^10^ Additionally, current environmental sample collection and sequencing methods recover predominantly short contigs. Short sequences are challenging to classify correctly because they often do not contain enough information (e.g., too few genes) to leverage our knowledge of what makes viral sequences distinct.^11,12^

The challenge of identifying viral sequences in metagenomic datasets has driven the development of many viral identification tools over the past decade that aim to differentiate viral sequences from non-viral sequences.^13^ Tools differ in the types of viruses they identify, what sequence lengths they are optimized for, and the training data and algorithms underlying them. To be confidently applied to environmental data, viral identification tools must be trained on sequences representative of the microbiota being studied to ensure the tool has seen enough of the sequence space to correctly classify viral sequences. Sequence types commonly found in environmental metagenomes include bacteria, viruses, plasmids, archaea, protists, and fungi. Some tools, such as VirSorter2^12^ and VIBRANT,^14^ include these diverse sequence types, as well as representative diversity within each sequence type, expanding the classification accuracy of each tool. Other tools like DeepVirFinder and VirSorter do not include as diverse sequences: DeepVirFinder does not include non-prokaryotic references and VirSorter is only built on bacterial and archaeal virus references. Further, the performance of viral identification tools depends on the interaction between the tool algorithm and the sample type. In a comparison of the viruses recovered by different viral identification tools across 13 environmental samples, differences were found in the number of sequences called viral between environments.^14^

While many viral identifications tools have comparable accuracy, their underlying algorithms differ and may capture different sets of viruses from the same sample. With so many tools available, it can be difficult to choose the most appropriate tool for a given study. Rather than choose one tool, a number of studies have combined the outputs of multiple tools to classify viral sequences to capture a greater portion of the viral signal.^15–21^ This approach assumes that combining multiple tools will improve overall accuracy by discovering more viruses without greatly increasing non-viral contamination (non-viral sequences called viral by the approach), but this assumption has not been rigorously evaluated. In particular, it remains unknown whether or not these multi-tool strategies significantly increase contamination (e.g., by each tool returning non-overlapping false positive viral sequences).

Here, we benchmarked whether multi-tool approaches can distill a more complete and accurate set of viral sequences. From our analysis, we recommend pipelines specific to short and long-read sequences, as well as cellular metagenomes and virus-enriched samples. By returning more viral sequences with less non-viral contamination, these pipelines will enable new and more accurate insights into the ways viruses impact microbial ecosystem functions, with far-reaching implications for human and environmental health.

## Methods

### Creation of sequence testing set

To create a testing set for benchmarking multi-tool pipelines, we downloaded viral, bacterial, fungal, plasmid, protist, and archaeal genome sequences from the NCBI reference sequence database (RefSeq), as well as a unique set of non-RefSeq virus genomes compiled by the VirSorter2 tool developers, herein ‘VirSorter2 database’^22^ (Figure 1A). These non-RefSeq virus genomes represent a comprehensive validated set of viruses. We originally created nine testing sets by randomly sampling sequences with replacement from these sources to create datasets that mimicked metagenomic environmental data. As the variability between testing sets plateaued at five datasets (Figure S1), five datasets were used for subsequent. The testing sets were designed to contain approximately 68% bacteria, 10% archaea, 10% virus (not proviruses), 5% plasmid, 5% protist, and 2% fungi sequences, totaling ∼8k sequences (Figure 1B). The proportion of sequences was chosen to be representative of cellular-enriched metagenomic data, which are dominated by bacterial sequences, and reportedly contain ∼10% viral sequences.^15,23^ The non-viral portion was randomly sampled from 5.3M bacteria, 55.1k archaea, 6.6k plasmid, 69.6k fungi, and 216.5k protist sequences from the NCBI database (accessed Nov 2019 for bacteria and archaea, April 2022 for others). The viral portion of the testing set was generated by random sampling from 13.8k viral sequences from the NCBI database and 370.153k viral sequences from the VirSorter2 database. As DeepVirFinder requires sequences to be less than 2,100 kb, a custom python script was written to trim the testing set sequences to meet this length cutoff. We did not expect trimming to impact results since the largest phage genome reported is smaller than this length cutoff (735 kb).^24^ Further, only sequences longer than 3 kb were used because it has been previously shown that tool accuracy significantly decreases below 3 kb.^12^

**Figure 1.**
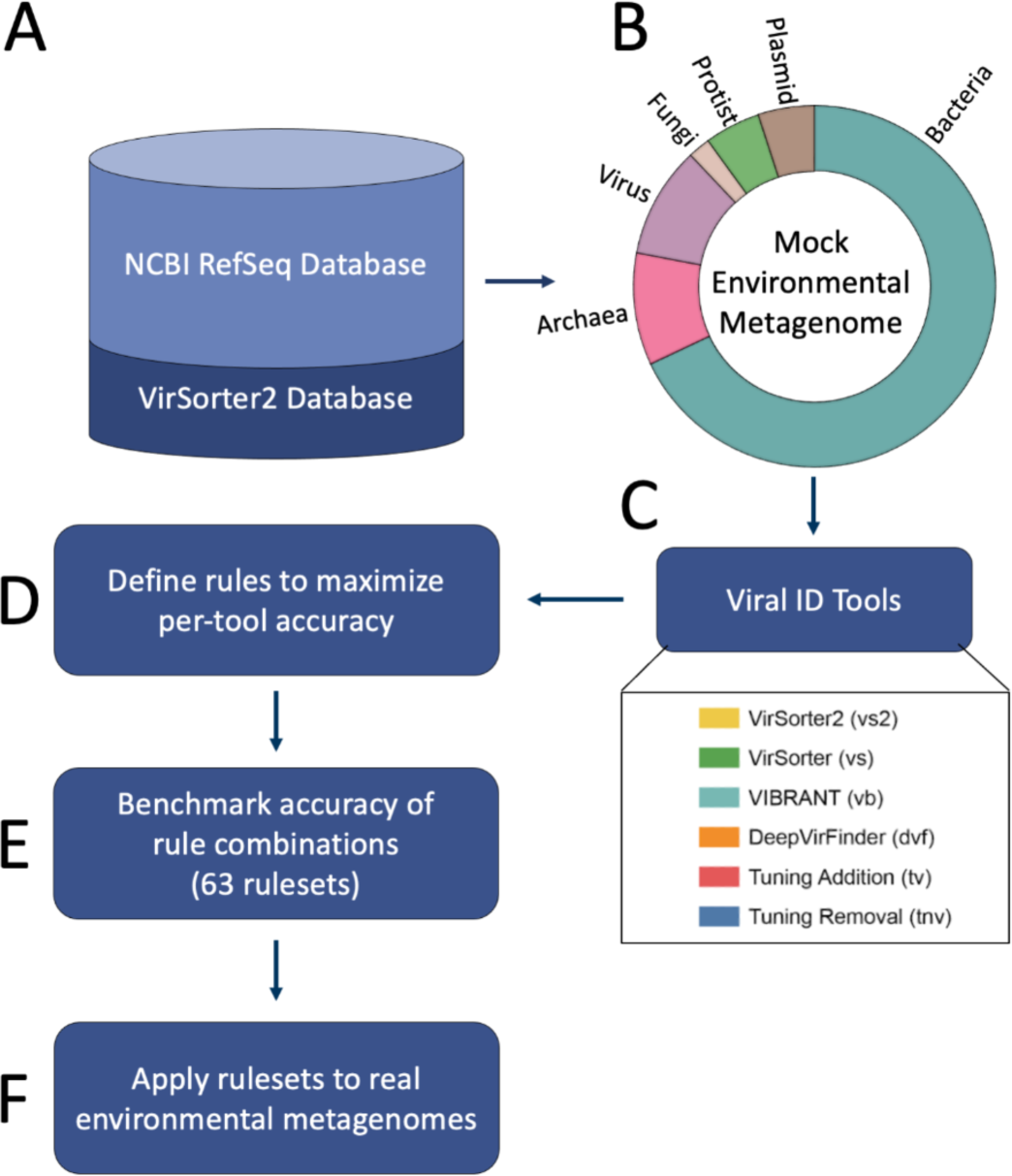
Overview of study workflow. (A) A set of sequences > 3 kb were randomly downloaded from NCBI and a curated Non-RefSeq viral genomes database (‘VirSorter2 database’) to (B) generate five mock environmental metagenomes, where the donut chart represents the proportion of each sequence type in each mock metagenome. (C) Mock metagenomes were run through six viral identification tools; (D) where score cutoffs were defined based on each tool’s outputs to maximize their accuracy. (E) Accuracy was then assessed for each tool combination to guide the development of defined “rulesets.” (F) Rulesets were then used to classify sequences from six real-world aquatic metagenomes: three cell-enriched metagenomes and three virus-enriched metagenomes.

### Selection of viral identification tools

27 viral identification tools^12,14,25,25–49^ were found through literature search and assessed to determine their suitability for inclusion in this study (Table S1). Tools were included if they met the following criteria: the tool (1) identifies viruses that infect prokaryotes (i.e., bacteria and archaea), (2) is suitable for application to multiple environments (e.g., not only the human gut), (3) is designed to target viral sequences of lengths greater than 3 kb, (4) can classify millions of sequences within a few days on high performance computing clusters (i.e., not only available on a web server), (5) developers actively respond to user issues, (6) performs well in previous comparative studies of viral identification tools,^12,14,50^ (7) is not specific to prophages, and (8) was published before June 2022.

Four of the 29 viral identification tools met the above criteria: DeepVirFinder,^27^ VIBRANT,^14^ VirSorter,^46^ and VirSorter2.^12^ While not designed strictly as viral identification tools, Kaiju^51^ and CheckV^26^ were used to tune the viral predictions in our test sets. All six are referred to as “viral identification tools” or simply “tools” in this manuscript (Table 1, Figure 1C).

**Table 1.** Overview of viral identification tools selected for inclusion in this study.

| Tool name (version) | Tool Description | Algorithmic Approach | Why we included the tool |
| --- | --- | --- | --- |
| CheckV<sup>26</sup> | CheckV is an automated pipeline that identifies closed viral genomes, estimates the completeness of genome fragments, and removes host regions from proviruses. | HMM virus and host marker genes, virus-host boundary prediction, AAI based estimation of genome completeness | Not a viral identification tool, but provides benchmarking information for refining predictions from other tools |
| DeepVirFinder<sup>27</sup> | DeepVirFinder uses a multi-layered deep learning algorithm trained on a positive set of viral sequences from viral RefSeq data and a negative set of prokaryotic ones. | K-mer based deep learning convolutional neural network | Recent and increasingly commonly used tool based on a neural network |
| VIBRANT<sup>14</sup> | VIBRANT is a hybrid tool that uses both machine learning and protein similarity to classify viruses as either high, medium, low quality, or non-viral. | Neural network of protein annotations of HMMs | Recent and commonly used tool, provides gene annotation information |
| VirSorter<sup>46</sup> | VirSorter uses probabilistic models with reference and non-reference dependencies, as well as detecting hallmark viral genes. | Probabilistic modeling using HMMs | Commonly used, high quality predictions |
| VirSorter2<sup>12</sup> | VirSorter2 uses a neural network classifier built on top of the existing VirSorter infrastructure of reference based viral identification. | A multi-classifier combining a Random Forest model and expert knowledge of viral features | Recent and increasingly commonly used tool that is more inclusive of viral diversity than most other tools |
| Kaiju<sup>51</sup> | Kaiju is a taxonomic classifier that compares metagenomic sequences to NCBI reference databases at the protein level and assigns a near or exact taxon match if one is found. | A taxonomic classifier that uses protein-level classifications to assign reads to taxon from NCBI databases | Not a viral identification tool, but extremely fast method of taxonomic identification based on NCBI releases |

The testing sets were run through each of the viral identification tools (CheckV (v0.9.0), DeepVirFinder (v1.0), Kaiju (v1.9.0), VIBRANT (v1.2.1), VirSorter (v1.0.6), and VirSorter2 (v2.2.3)) using the University of Michigan Great Lakes Supercomputing Cluster. The Kaiju nr_euk database (updated from NCBI 05-23-2022) was used for Kaiju taxonomic classification. All tools were run with default parameters except for specifying a 3 kb contig length cutoff.

### Design of viral identification rules

Viral identification tools generate scores that indicate how confident they are that a given sequence is of viral origin, but users are often faced with the dilemma of setting their own score cutoff to decide which sequences to call “viral.” To aid in the process of choosing rules and cutoffs for predicting viral sequences, we designed six rules (Figure 1D, Figure 2). Each rule includes at least two *subrules* that use outputs from the six selected tools. These subrules were designed through two processes:

**Figure 2.**
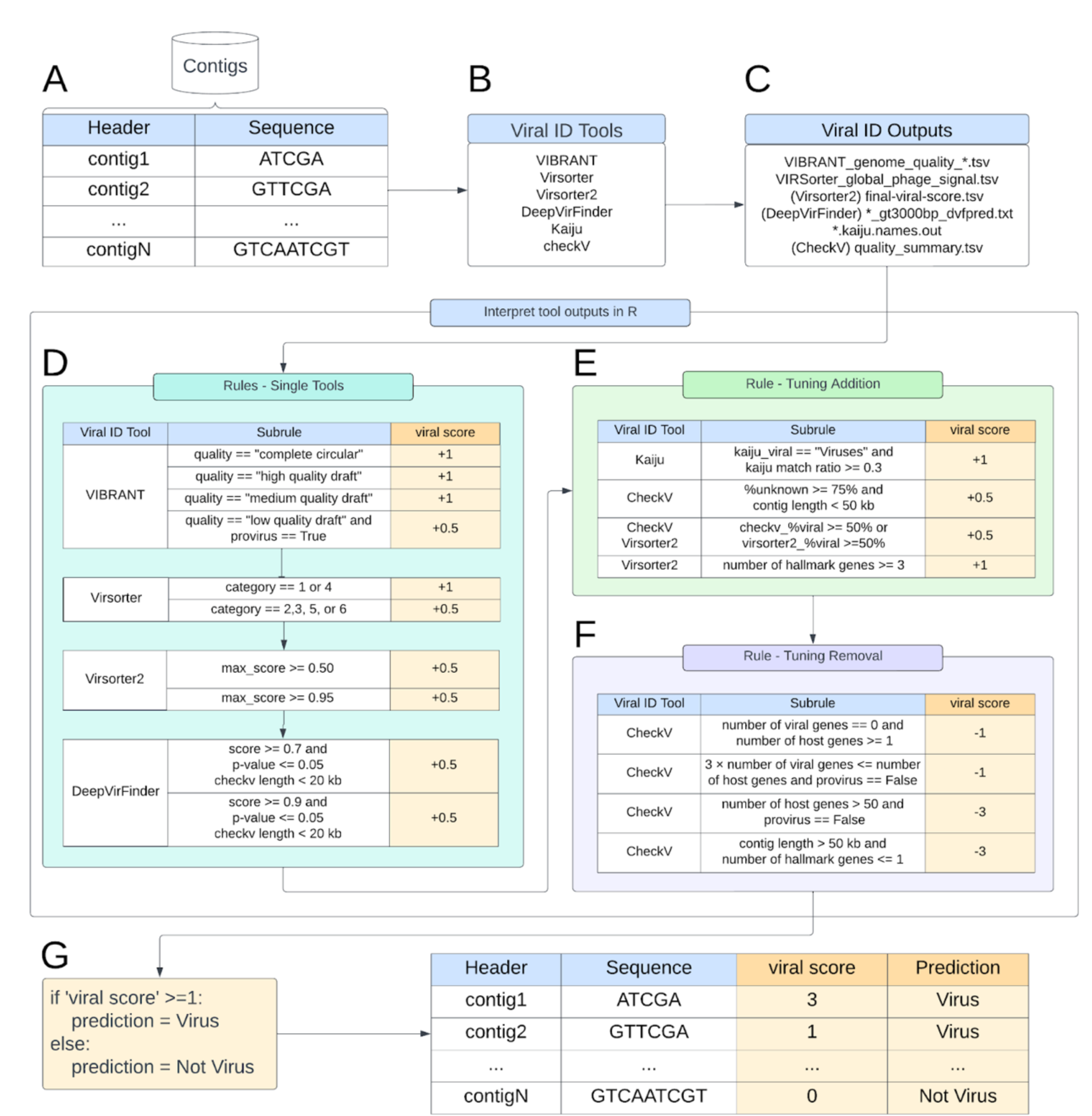
Diagram of approach details. (A) Sequences are first processed by each (B) viral identification tool. Next, (C) the tool outputs are programmatically post-processed to generate a *viral score* based on both (D) single-tool rules and the data-driven creation of (E) tuning addition and (F) tuning removal processes. (G) This combined post-processing generates a *viral score* that indicates whether each sequence input is predicted as “Virus” or “Not Virus.” Subrules are scored based on the confidence of the prediction: low-confidence = ±0.5, confident = ±1, highly confident not viral = -3.

(1) *Evaluation of existing recommendations* for *tool cutoffs and application*: the recommended cutoffs for distinguishing viral and non-viral sequences in each tool’s protocol were used as an initial set of rules.^12,15,16,52^

(2) *Curation and evaluation of biological* features: some viral identification tools generate information describing biological features for each sequence, e.g., VirSorter2 reports the number of viral hallmark genes identified, CheckV reports the completeness of a sequence and relative percentage of viral versus non-viral genes, VIBRANT identifies virus orthologous genes (VOGs). These biological features were used to create classification criteria (Figure 2) to distinguish viral and non-viral sequences.

The developers’ recommendations for calling a sequence viral served as a baseline for assigning a given sequence a *‘viral score’* that captured the relative likelihood that a given sequence was viral. The cutoffs were then adjusted to maximize the number of true viral sequences being classified as viral (true positives) and minimize the number of non-viral sequences being classified as viral (false positives) in the mock environmental microbial communities. We first defined four single-tool rules (Figure 2A-D) derived from four viral identification tools (i.e., VIBRANT, VirSorter, VirSorter2, DeepVirFinder). Next, we defined two sets of tuning rules derived from outputs of different tools that indicated a strongly viral or non-viral signal (Figure 2E-F). These included: (1) a “tuning removal” rule that decreases the *viral score* based on distinctly non-viral sequence features and (2) a “tuning addition” rule that increases the *viral score* based on distinctly viral sequence features.

Each putative viral sequence is assigned a numerical value by each rule; these are combined to give a value that comprises the sequence’s *viral score* (Figure 2G). Sequences with a final *viral score* ≥ 1 were considered viral, and scores < 1 non-viral (Figure 2G).

### Evaluation of viral identification rulesets

Ultimately, 63 combinations (rulesets) of these six rules were evaluated by comparing the viral score of each sequence to the classification assigned by the database (Figure 1E). From these values, precision (the number of true viruses in our test set called viral divided by all contigs called viral), recall (the number of true viruses in our test set called viral divided by all true viruses), and Matthews Correlation Coefficient (MCC, considers the relative proportion of false positives, false negatives, true positives, and true negatives)^53^ were calculated.

### Application of rulesets to environmental metagenomes

All rulesets were used to identify viruses from five previously published environmental datasets representing different aquatic environments and size fractions (Figure 1F, Table S2). Three environments (drinking water, global ocean water, and eutrophic lake water) contained metagenomic assemblies (>2 µm), and three environments (wastewater, eutrophic lake water, oligotrophic lake water) contained virome assemblies (<0.2 and <0.45 µm), meaning samples were enriched for viruses by filtering through a small pore to remove most cellular organisms before DNA extraction.

The default tool settings were used except for the oligotrophic lake and wastewater virus-enriched samples, where the -viromè flag was used for VIBRANT and VirSorter to reduce sensitivity, as recommended by the developers of those tools given that a greater fraction of total sequences are expected to be viral in virus-enriched samples. The ‘-virome’ flag was not used for the eutrophic lake water virome assemblies due to all eutrophic lake water assemblies being processed together, but the eutrophic lake water virome still was most similar to the other virus-enriched samples (Figure 7). All tools were run with a 3 kb cutoff to remove small sequences.

### Code availability

The scripts to run the full testing pipeline presented above, including those to identify viral contigs from these tool outputs, example runs, and classifier outputs on the environmental metagenomic samples, are freely available at https://github.com/DuhaimeLab/VSTE.

## Results and Discussion

Viral identification tools use algorithms based on knowledge of viral sequences and machine learning to separate viral and non-viral sequences. In this study, we test the hypothesis that combining viral identification tools with different underlying algorithms will improve accuracy. Performance of 63 combinations (rulesets) of the six rules were evaluated using five mock metagenomes of known composition (Figure 3B, S1). The six rules are as follows: four single-tool rules derived from four viral identification tools (i.e., VIBRANT, VirSorter, VirSorter2, DeepVirFinder; Figure 2A-D) and two additional tuning rules: tuning removal (which removes predictions using Kaiju, CheckV, VirSorter2, VirSorter, and VIBRANT outputs) and tuning addition (which adds predictions using Kaiju, CheckV, and VirSorter2 outputs; Figure 2E-F). In this section, we compare the performance of our rulesets, elaborate on their strengths and weaknesses, and provide recommendations for use.

**Figure 3.**
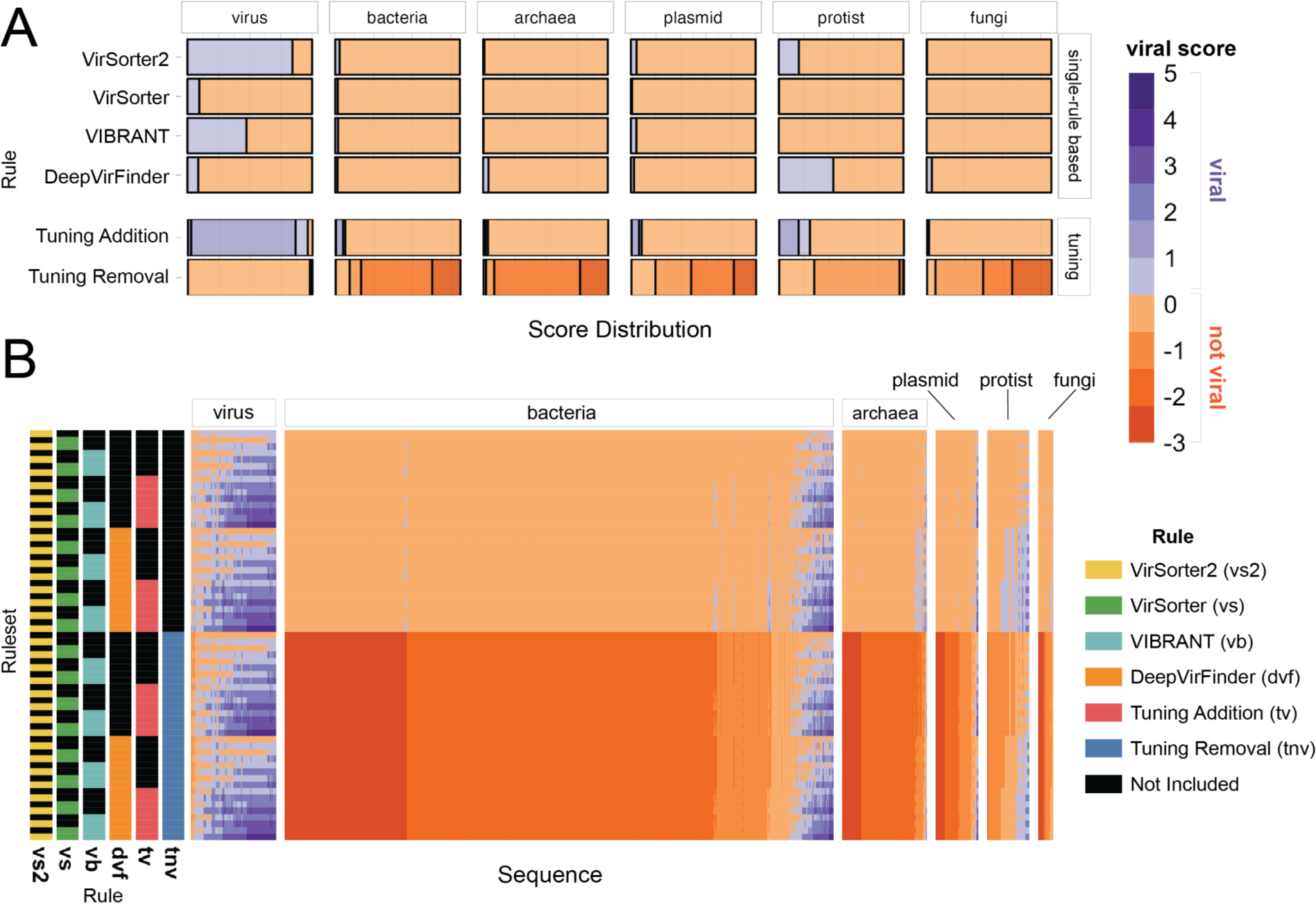
Comparison of different rulesets. (A) Distribution of *viral scores* assigned to mock metagenome sequences for our 6 rules: 4 single-tool and 2 tuning rules. (B) Viral scores of all sequences across six mock metagenomes classified by each ruleset. Ruleset rows are colored based on whether or not each rule was used to attain the *viral score* result for a given sequence. In both A and B, *viral scores* ≥ 1 are classified as viral; and < 1 as not viral. All sequences are grouped by their assigned taxonomy.

### More tools are better… to a point

Across the 63 rulesets, MCC, our metric for overall performance (‘accuracy’, herein), ranges from 0.05 (DeepVirFinder) to 0.77 (VirSorter2 + Tuning Removal). With the exception of VirSorter2 (MCC=0.75), single-rule rulesets (i.e., viral identification tools run on their own) either missed most of the viruses in the benchmarking dataset or misclassified such a large number of non-viruses that the viral signal was heavily contaminated (Figure 3A). Of the single-rule rulesets, VIBRANT performs second best (MCC of 0.55), followed by VirSorter (MCC of 0.16) and DeepVirFinder (MCC of 0.05). Previous studies have reported higher accuracy for these tools, but those studies used a testing set composed of 50% or more viruses compared to our 10% viral sequences and/or did not include taxonomically diverse sequences compared to the taxonomically diverse training data used here.^14,27,46,52^ Using testing datasets representative of environmental metagenomic datasets is important. We found that our MCCs increased when using a testing set with a more similar composition to other studies (50% viral and 50% non-viral; max MCC=0.91); and, in their validation of DeepVirFinder, Ren et al.^27^ demonstrated that the relative proportion of viral to non-viral sequences in a dataset can have a strong effect on AUROC (area under receiver operating characteristic; a performance metric). Given the observed taxonomic distribution of environmental cellular metagenomic sequences,^15,23^ users likely need to be more conservative (higher classification score cutoffs) in viral calling and assume lower accuracy than previous studies have reported.

Combining rules generally increased the average MCC (Figure 4). This is driven by a statistically significant increase in recall for multi-rule rulesets compared to single-rule rulesets and for three or more rule rulesets compared to two-rule rulesets (Table S3). While average precision is constant as rules are combined (Figure 4A, Table S4), the precision of higher precision rules (precision > 0.7, five right-most points Figure S2A; the VIBRANT and VirSorter2 rules) decreases as they are combined with lower precision rules (precision < 0.5; the DeepVirFinder and VirSorter rules). Accuracy is maximized by the VirSorter2 and tuning removal ruleset, and is not improved by adding more rules (Figure 4B).

**Figure 4.**
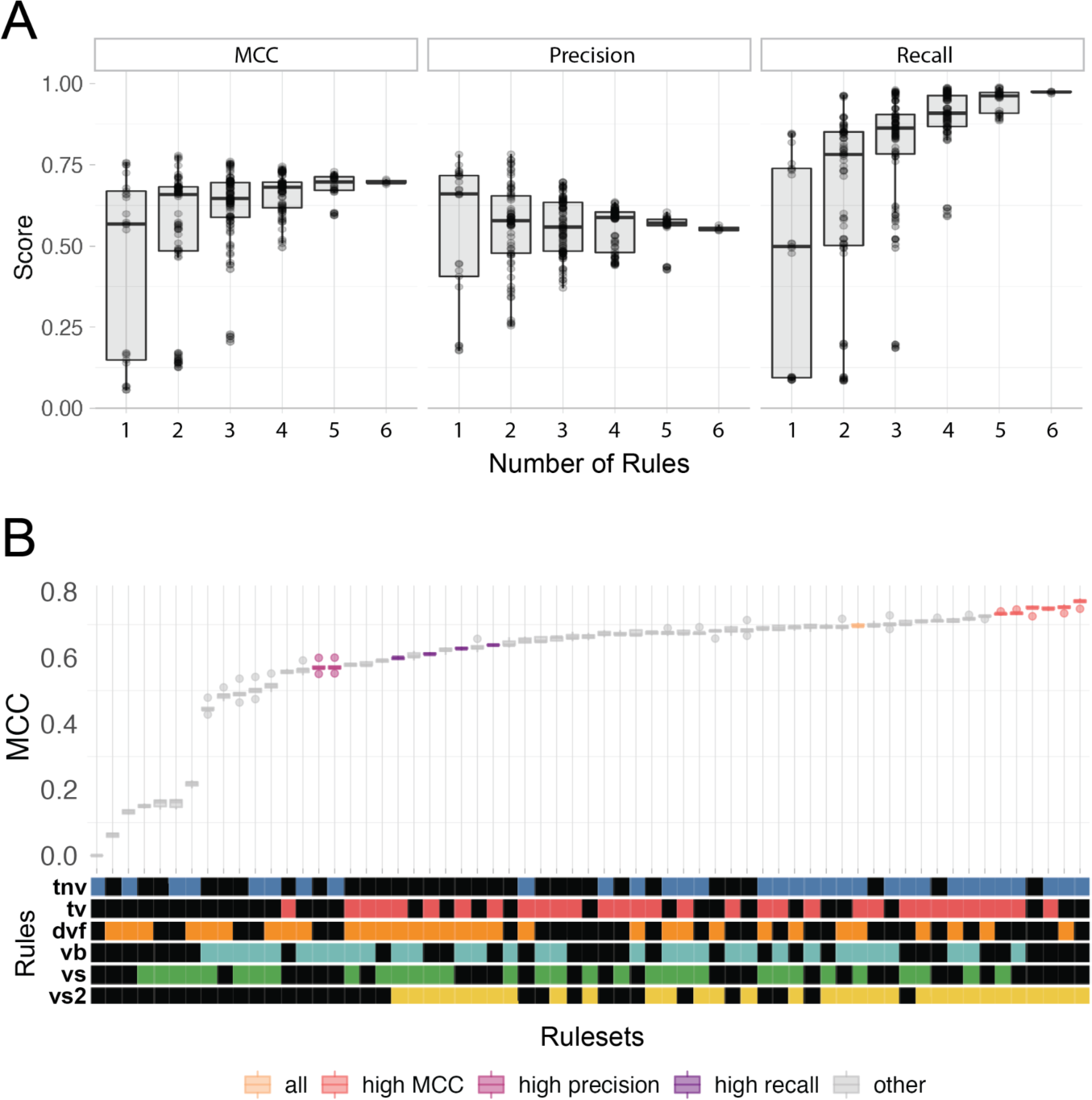
Performance of the 63 rulesets. (A) - Box and whisker plots of the performance scores representing variation in MCC, precision, and recall of different rulesets based on the number of rules used for prediction. (B) Ruleset accuracy (MCC) ordered by increasing MCC and colored based on the ruleset’s type according to statistically equivalent (p_adj_ ≥ 0.05) rulesets. For A and B, the middle line represents the group mean; boxes above and below the middle line represent the top and bottom quartiles, respectively; whiskers above and below the boxes represent 1.5 times the interquartile range (roughly the 95% confidence interval), outliers are represented by circles beyond the whiskers. The boxplots in A are overlaid with points that represent each testing set’s MCC.

The VirSorter2-based and tuning removal rules are the most critical for accurate virus identification in our testing. When rulesets were ranked by increasing MCC, top performing “high MCC” rulesets were identified as those that did not demonstrate a statistically significant decrease relative to the highest MCC ruleset (p_adj_ > 0.05; Figure 4B). VirSorter2’s rule (‘vs2’) was in all six of these high MCC rulesets (Figure 4B) and tuning removal (‘tnv’) was in five of them. For comparison, the other single-tool rules (i.e., VirSorter, DeepVirFinder, and VIBRANT) were each only in one of the high MCC rulesets, where they always co-occurred with VirSorter2 and the tuning removal rule (Figure 4B). None of the high MCC rulesets have more than four rules. In the same way, “high precision” (three rulesets; Fig S2A) and “high recall” (four rulesets; Fig S2B) rulesets were defined.

There is a high degree of overlap in the viruses predicted by the different rulesets (Figure 5). 68% of the comparisons between rulesets (Figure 5, green and yellow cells) were more than 50% identical to each other; and 30 rulesets were at least 90% identical to at least one other ruleset (Figure 5, yellow cells), representing 4% of the total comparisons between rulesets. Rulesets with VirSorter, DeepVirFinder, and VIBRANT all have more sequences in common as the number of rules in the compared sets increases, but this trend is much less pronounced for VirSorter2 (Figure S3). This suggests that our observed increase in recall through combining rules is being driven by a subset of rules that give more tool-specific viral predictions (i.e., VirSorter, DeepVirFinder, and VIBRANT), leading to the question: “how confident should users be of their tool-specific predictions?”.

**Figure 5.**
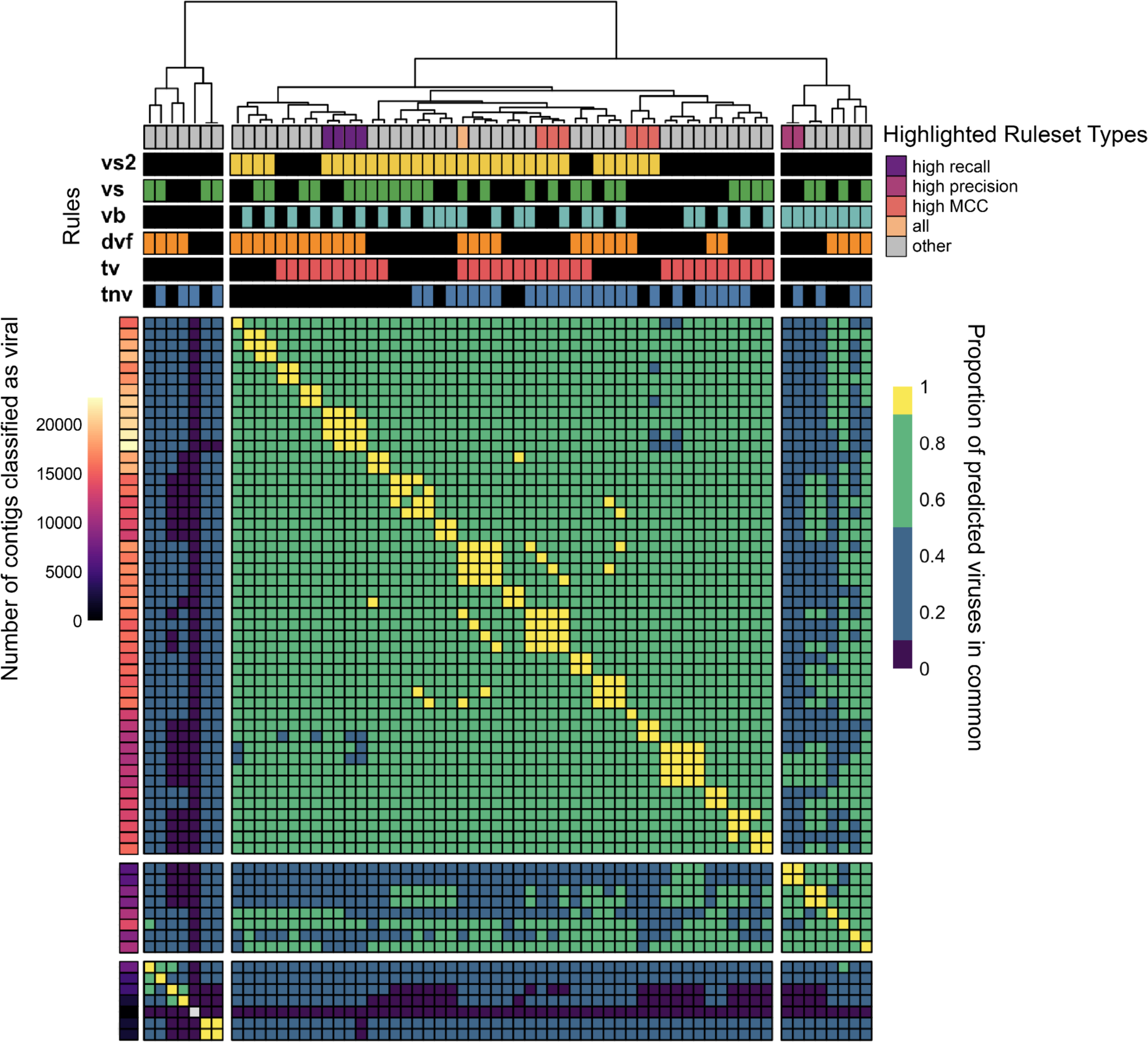
Proportion of viruses in common between rulesets. Heatmap values calculated by dividing the intersection (called viral by both rulesets) by the union (called viral by at least one) of the viruses found by both rulesets, which represents the proportion in common between the tools (scale bar on right: dark purple: 0-0.1, blue: 0.1-0.5, green: 0.5-0.9, yellow: 0.9-1). The bar to the left of the heatmap represents the total number of viruses identified by each tool combination (scalebar to its left). The bars above the heatmap indicate the tool(s) used in the rulesets, as well as the ruleset type.

One assumption made during *in silico* virus identification by previous studies is that sequences with low-confidence predictions by multiple tools are more likely to be viral (i.e., called viral by multiple +0.5 subrules). For example, if one tool predicts a sequence as viral with low confidence, it may be disregarded. But if multiple tools each provide low confidence predictions for a given sequence, many studies have presumed the sequence is more likely to be viral, thereby combining low quality predictions to arrive at the set of predicted viruses.^15,16,18^ However, we found this was not a safe strategy as it did not increase the number of true-positives. Rulesets using multiple low-quality prediction subrules in the single-tool rules did not significantly increase MCC (p-value=0.19) or recall (p-value=0.18); and in fact, slightly decreased precision (0.54 vs 0.59, p-value=2.5*10^-5^) when compared to rulesets that did not use low-quality prediction subrules (Figure S4). This pattern is likely due to the rulesets being similarly uncertain about sequences that are unlikely to be viral; thus, introducing a significant amount of non-viral contamination. The additive uncertainty did not create certainty. We recommend being cautious of sequences classified as viral by multiple low-quality predictions and manually inspecting them before viral assignment.

As metagenomes frequently have a high proportion of short sequence fragments and the correct identification of short fragments with only a few genes is particularly challenging,^26^ we tested the rulesets on 3-5 kb fragments from our original testing sets. Like the >3 kb testing sets, many rulesets performed similarly for the 3-5 kb fragments (Figure S6). Unlike in the >3 kb testing sets where the VirSorter2 single-rule ruleset was in the high MCC set (Figure 4B), no single-rule ruleset was in the high MCC rulesets for the short fragments (Figure S6). Also in contrast with the >3 kb testing sets, the six-rule ruleset was identified as a high MCC ruleset and DeepVirFinder is in 8 of the 12 high MCC rulesets. In part due to our newly defined viral tuning rules, the accuracy of the viral predictions reported here for these short fragments is greater than previously published.^12^ This increase in accuracy suggests that datasets where short fragments abound may particularly benefit from our tuning rules. Further, while researchers with only 3-5 kb fragments may consider using more tools, equally accurate predictions can still be achieved from the VirSorter2 and tuning removal ruleset.

### Tuning rules increase confidence of viral predictions

To leverage expert knowledge of the differences between viral and non-viral sequences, we designed tuning addition and removal rules (Figure 2E-F; see Figure S6 for subrule performance). These rules were designed based on specific outputs from multiple tools that distinguish between viral and non-viral sequences, such as sequence length and the number of host genes. In general, tuning addition improved MCC and recall, while tuning removal improved MCC and precision (Figure S7). 7 of the 10 highest MCC rulesets have both tuning addition and removal rules (Figure 4B), demonstrating the importance of the tuning rulesets for accurate classification. The tuning *removal* rule was able to identify 89% of the testing set’s non-viral sequences and only (mis)identified 2% of testing set’s viral sequences as non-viral (*viral score* < 0 when only the tuning removal rule was applied). The tuning *addition* rule accurately identified 74% of the viral sequences and only misidentified 4% of non-viral sequences in the testing set as viral (*viral score* ≥ 1 when only the tuning addition rule was applied). Overall, the tuning rules increased our prediction accuracy beyond that of the rulesets composed of the single-tool rules. As such, we demonstrated the value of automating the refinement of viral identification tool predictions, a task that, if done at all, is currently a laborious manual process.

Even with the tuning rules, we could not improve both precision and recall beyond 0.77 (Figure S8). Building a more accurate classifier means overcoming barriers such as imperfect gene reference datasets, overlap between viral and host sequences, and underrepresentation of viral types. This is because to recover more viruses, it becomes necessary to rely more on genes of unknown origin. These may include non-viral genes, particularly of eukaryotes, which were not represented in the reference datasets of DeepVirFinder, VirSorter or CheckV (Figure S9-16). Further, many true viral features overlap with non-viral features (Figure S9-16) due to our imperfect knowledge of what distinguishes viruses and non-viruses (and homologous sequences shared by both viruses and cellular organisms), leading non-viruses with virus-like features to be misclassified as viruses. This challenge is particularly acute when trying to accurately classify both short sequences (<5 kb)^12,27^ and viral types underrepresented in our testing data (e.g., the accuracy of the “high MCC” ruleset is highest for dsDNAphages compared to other viral sequence types, Figure S17).

### Mislabeled sequences within databases hinder tool accuracy and validation

To improve upon the maximum MCC of 0.77, we looked for patterns in the types of sequences being misclassified that could aid future tool design. To our surprise, the “false positive” sequences labeled as bacteria by the NCBI database, but classified as viral by our high MCC ruleset, looked more “viral” than the viruses themselves. Specifically, the proportion of the sequence’s genes identified as viral orthologous genes (VOGs) was higher in the misclassified bacteria than the known viruses (p-value < 2.2*10^-16^; Figure 6A). The plasmids misclassified as viral have a similar proportion of viral genes as the viruses (p-value = 0.63). For all three sequence types, the proportion of sequences represented by VOGs increases with the number of VOGs in that sequence (Figure 6B). If these highly viral sequences are not actually viruses, viral identification tools can be improved by removing these genes from viral gene databases. On the contrary, if these sequences are actually viruses, viral identification tools can be improved by relabeling these sequences in sequence databases because tools rely on accurate database classification for training and testing.

**Figure 6.**
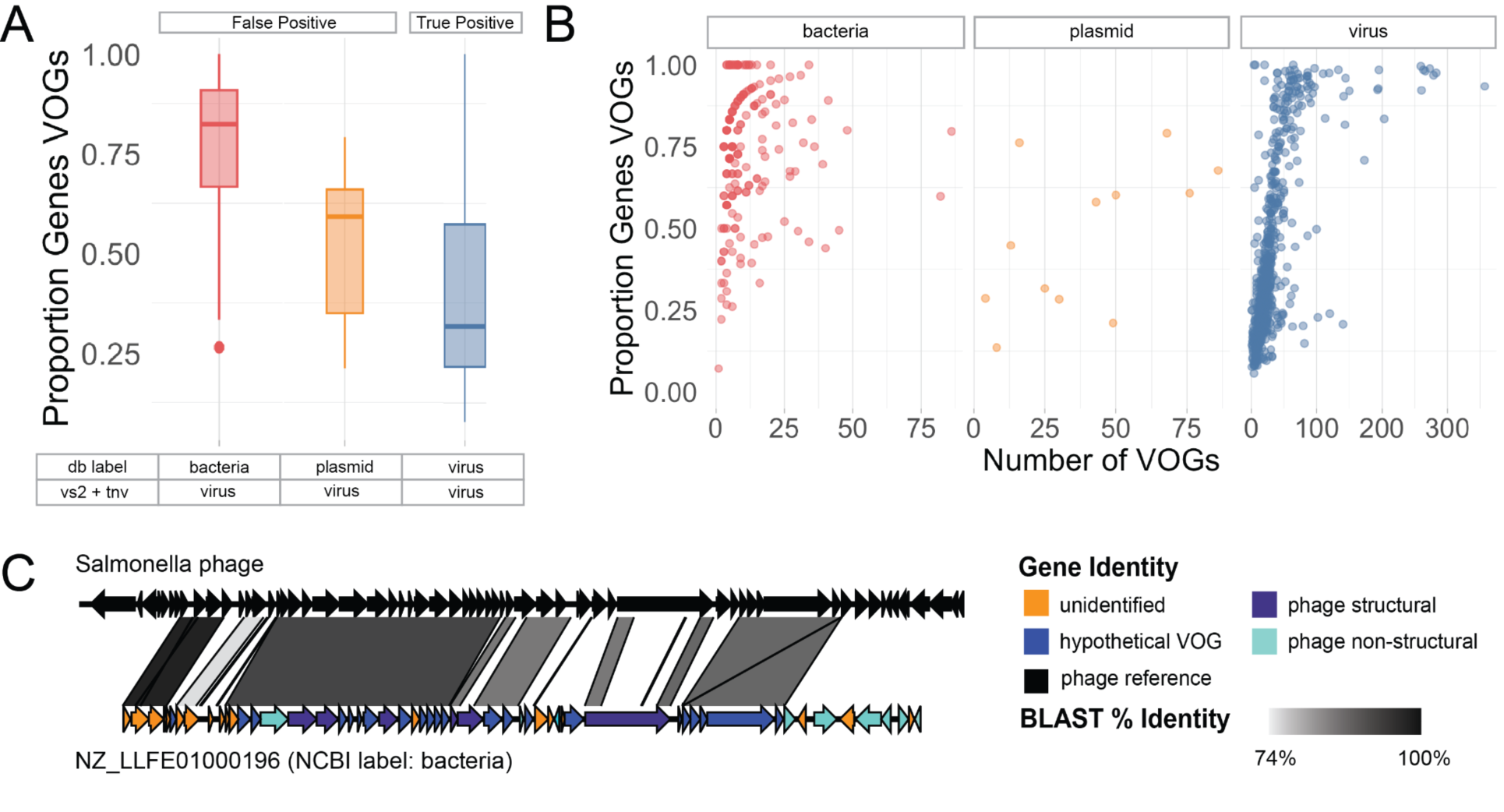
Mislabeled sequences. (A) Box and whisker plot of the proportion of genes on a sequence with a VOG annotation by VIBRANT broken down by sequence type (for the high MCC rules). (B) Proportion of a sequence’s genes with a VOG annotation versus the number of genes with a VOG annotation faceted by sequence type. Because VIBRANT is the only tool that provided VOG annotation, only sequences classified as viral by VIBRANT are included in panels A and B (which only included bacteria, viruses, and plasmids). (C) Sequence synteny plots indicating sequence similarity between a representative bacterial “false positive”: NZ_LLFE01000196 (NCBI label: bacteria) versus Salmonella phage SSU5. All genes of the testing set sequences are colored by their gene identity.

Manual inspection of a subset of these misclassified sequences revealed them to be viral sequences (Figure 6C, S18). These false positives represented two types of mislabeled sequences: (1) viruses (either extracellular virions or intracellular extrachromosomal viral genomes) co-sequenced when a host isolate was sequenced (Figure 6C and S18A) and (2) prophages integrated into their hosts’ genome (Figure S18B). We also screened for ΦX contamination, as it is a known problem for NCBI database sequences using Illumina library preparation due to its use in Illumina libraries.^54^ Only 19 ΦX sequences were taxonomically identified by Kaiju; and, thus do not explain our high degree of false positives. These findings support the known problem of phage sequences not being removed before being deposited on NCBI,^54^ and lead to phages being misclassified as bacteria, archaea, plasmids, and cellular and satellite chromosomes. Mislabeled sequences in public databases make it difficult to produce accurate viral identification tools because developers rely on accurately labeled datasets to train and test their classifiers. We recommend that before uploading sequences to public databases that non-viral sequences are screened for viral contamination.

### Sample preparation and viral identification tool choice affects viral sequence recovery

To compare our rulesets across environments, we evaluated the proportion of sequences classified as viral for five publicly available aquatic metagenomic assemblies (Table S2, Figure S19). We found that viral sequence recovery varied greatly based on sample preparation (e.g., virus-enriched or not) and viral identification tool(s) used (Figure 7). The highest MCC ruleset identified a higher proportion of viral sequences in the virus-enriched samples (44-46%) compared to the non-enriched metagenomes (7-19%). The proportion of viruses recovered across rulesets for the environmental datasets mimics the behavior of the testing data: the “high recall” rulesets classified the most sequences as viral (with presumably the greatest non-viral contamination, given the results on our testing sets), while the “high precision” rulesets labeled the fewest sequences as viral (with presumably the least non-viral contamination, given the results on the testing sets) (Figure 7, and Table S2). Previous metagenomic studies of biomass collected on a 0.2 µm filter have found viral sequences to be less than 16% of the total metagenome,^15,23^ which is in line with our results. Further, these ranges in cellular-enriched samples (>0.2 µm fraction) is consistent with the ∼10% that one would expect, given that viral genomes tend to be 1-2 orders of magnitude smaller than their microbial hosts and are on average an order of magnitude more abundant.^15^

**Figure 7.**
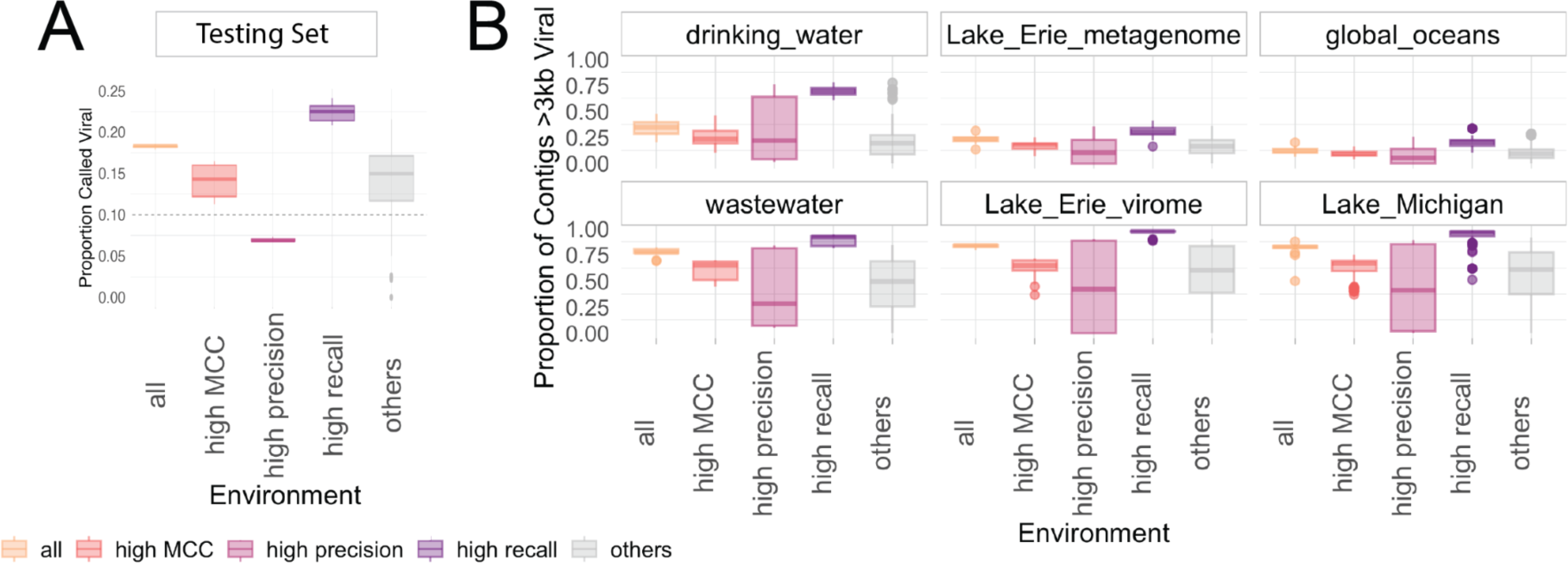
Proportion of viruses predicted by each tool combination across. (A) our testing sets and (B) environmental datasets. Rulesets are grouped based on the accuracy type on the testing set shown in the highlighted rulesets from Figure 5.

For the virus-enriched samples, nearly all sequences were called viral by the “high recall” rulesets (Figure 7, S18). It is likely that some of these sequences are false positives because the tuning removal rule reduces the number of sequences called viral by 14.9% (1 SD = 2.1%) when comparing the “all” to the “highest recall” rulesets; and our benchmarking demonstrated that the tuning removal rule effectively removed the contaminating non-viral sequences without removing true viruses. Further, one of the few studies to report the proportion of viral sequences found that 30-60% of the sequences with known taxonomy were similar to at least one viral sequence.^55^ Even if all unknown sequences were viruses in this study, the proportion of viruses would not exceed 92%, which is still lower than the proportions we found in two of our three virus-enriched samples; and, further suggests the importance of tuning removal even for viral-enriched metagenomes. We present this “high recall” example to caution readers against using the “high recall” rulesets on virus-enriched metagenomes. It may be tempting to assume virus-enrichment removes nearly all non-viral sequence contamination, but even for virus-enriched fragments we instead recommend using the tuning removal rule unless the number of sequences is small enough to be manually inspected.

### Recommendations and Future Work

*In silico* prediction of viral sequences is a critical first step to any metagenomic study that aims to resolve viral ecology and virus-microbe interactions. As downstream analyses and conclusions are predicated on accurate viral prediction, it is paramount to choose the most suitable tool(s) (and know how to interpret its output(s)) among the rapidly increasing number of *in silico* viral prediction tools available. Through the above benchmarking, we demonstrated that specific two-rule rulesets provide the highest precision viral identification with only minor sacrifices in recall, whereas the worse performance of combining all tools may lead to erroneous biological conclusions. For this reason, we urge caution when using recent automated pipelines for sequence identification that combine the output of multiple tools.^17,19–21^

Our recommendations based on this study vary depending on research question and experimental design (Figure 8). For a typical study investigating viral diversity and functional potential from a mixed metagenome, we recommend our “high MCC” ruleset (VirSorter2 with tuning removal). If the majority of eukaryotes were filtered out of the sample (e.g., <3 µm fraction was sequenced), the tuning additional rule may increase recall. For researchers seeking to minimize the number of tools they use, we recommend using VirSorter2 with our CheckV-based tuning subrules. VirSorter2 has a comparable MCC to multitool rules (Figure 4), though its high recall comes at the cost of more false positives when used in isolation (Figure S2). While VIBRANT’s high precision with and without our tuning removal rule (Figure 4, Figure S2), as well as convenient information about the viruses (e.g., their metabolic potential), are attractive, we found that its recall was much worse than our other recommendations on the environmental datasets. In general, however, we do not recommend researchers to use any of the tools in isolation, based on the poor accuracy of the single-rule sets on the short sequences (3-5 kb testing set, Figure S5).

**Figure 8.**
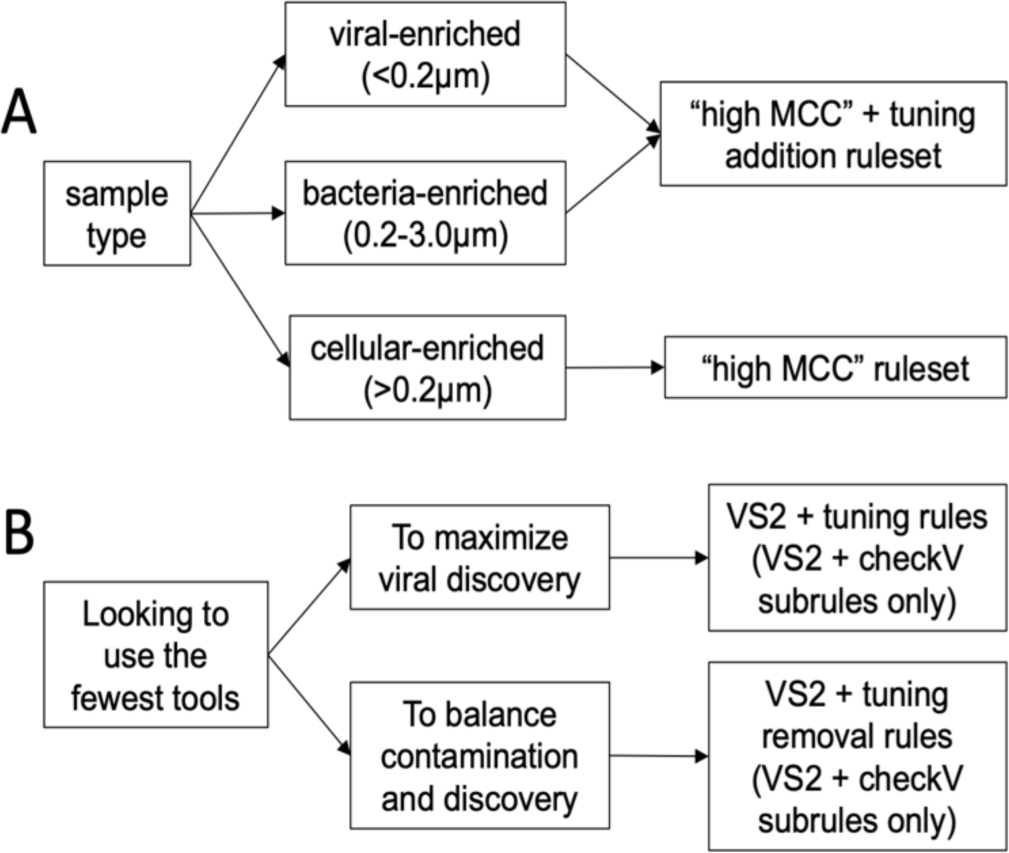
Recommendations. Flowcharts are based on (A) sample type and (B) study goals when looking to minimize the number of tools used.

One limitation of this work is that the available testing data included sequences that were part of the tools’ original training and testing data: all tools included in this study were trained in part using NCBI sequences that overlap with our testing data, and some were trained using the VirSorter2 non-RefSeq genome set. These datasets also happen to harbor a large number of mislabeled sequences, making it difficult to assess the accuracy of the benchmarking results. As additional curated viral sets are published,^56^ other researchers can test our rules against the new datasets providing further information about the limitations and scope of our rules.

We focused on tools that were developed primarily for bacteriophage identification, as these are most commonly used by microbial ecologists and microbiome researchers. We did not evaluate tools that were specifically for prophages, human viral pathogens, eukaryotic viruses, or archaeal viruses more broadly (Table S1). Future integration of new tools for plasmid and eukaryotic sequence identification^54,57^ is likely to improve viral identification tool pipelines.

## Conclusions

With the rapid development of new viral identification tools, this paper offers a blueprint for intentional, data-driven validation of tool combinations. We found that the highest accuracy resulted from rulesets with four or fewer rules. For most applications, we recommend a combination of VirSorter2 and tuning rules based on features of viral and non-viral sequences; and caution against simply combining viral identification tools expecting higher quality virus sets. By increasing the proportion of high-confidence viruses identified from mixed metagenomic datasets through intentional, data-driven combination of tools, this study enables more accurate ecological analyses by decreasing contamination of the viral signal, particularly from eukaryotic sequences.

## Acknowledgements

This work was supported by the Blue Sky Initiative of the University of Michigan College of Engineering (B.H., M.B.D.), National Science Foundation award #2055455 (M.B.D., E.B., M.L.), the National Science Foundation Graduate Research Fellowship Program under Grant No. DGE-134012 (J.R.) and DGE-1256260 (M.L.), and the National Oceanic and Atmospheric Administration Great Lakes Omics program distributed through the UM Cooperative Institute for Great Lakes Research NA17OAR4320152 (M.B.D, A.W.).

