## Supplementary material for "Benchmarking informatics approaches for virus discovery: Caution is needed when combining *in silico* identification methods": SI Figures

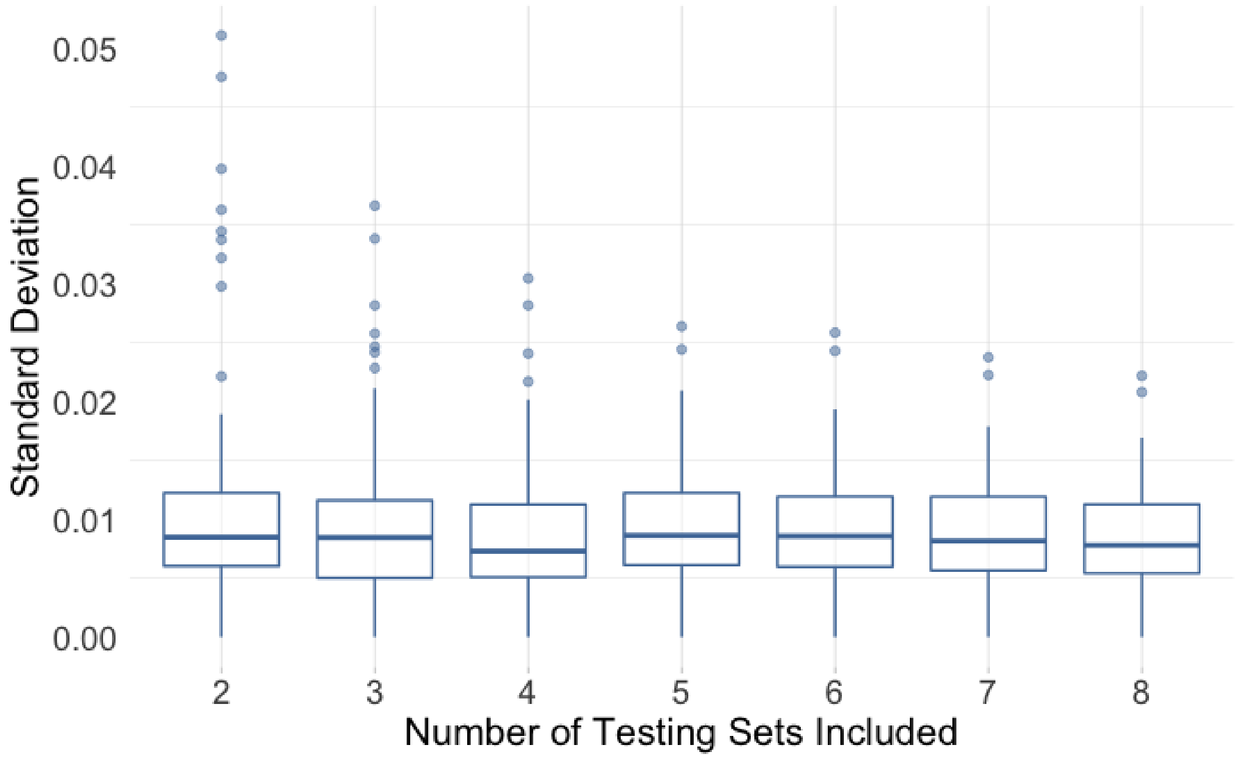


**Figure S1**. Variability across testing sets. The average standard deviation is consistent across the number of testing sets (p-values $\geq 0.25$), and the variability of the standard deviation plateaus after 5 testing sets are included (p-values $\geq 0.65$).

A

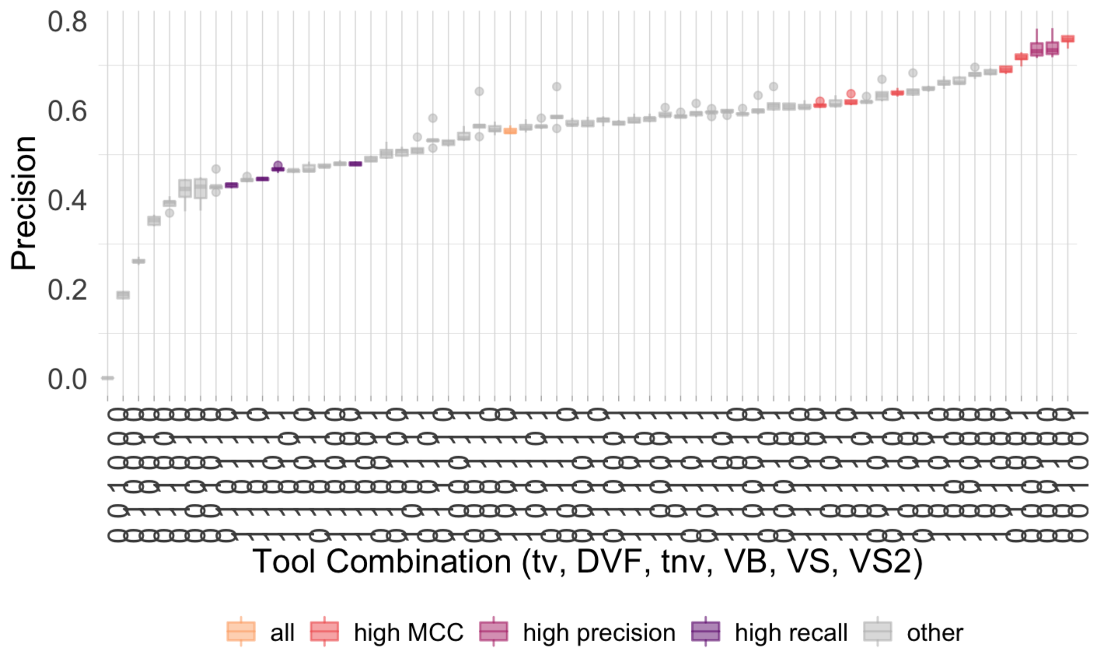


B

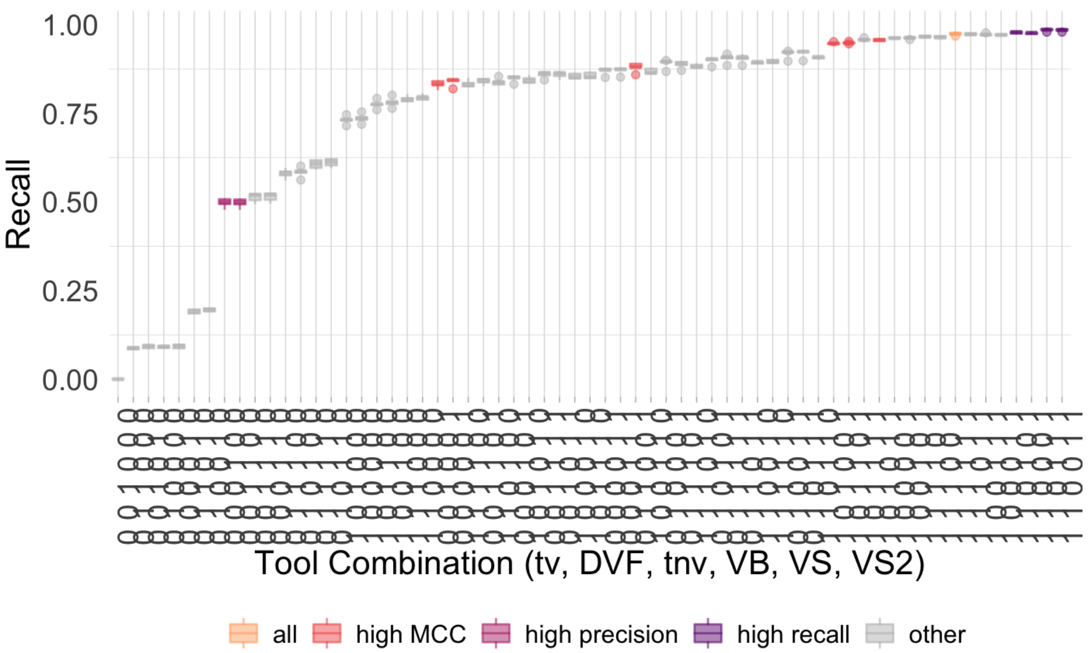


**Figure S2**. (A) Precision and (B) recall boxplots by ruleset. Ordered by increasing precision and recall, respectively. Colored by the ruleset type.

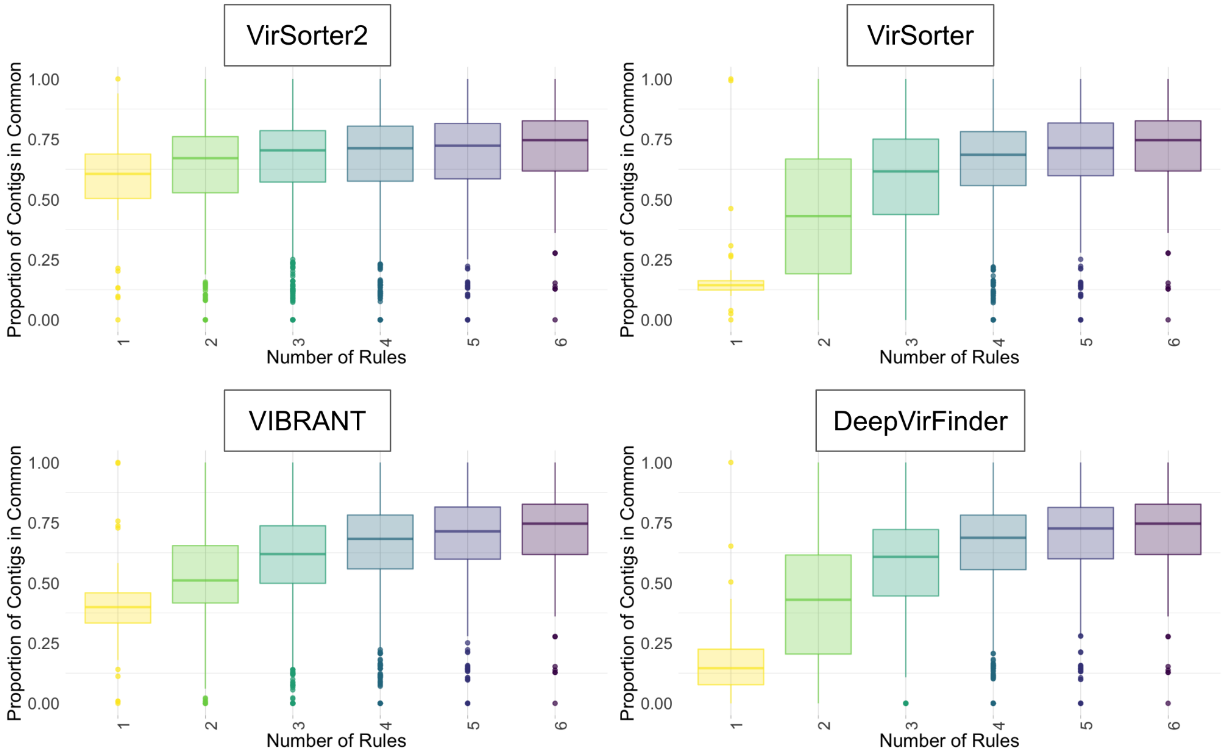


**Figure S3**. Proportion of sequences in common between two ruleset combinations for the four viral identification tools tested (VirSorter2, VirSorter, VIBRANT, DeepVirFinder). The combinations are grouped along the x-axis by the number of rules in the first member of the pair. For this reason, a combination with a ruleset with 1 rule and a ruleset with 3 rules would be included once in the "1" rule column and once in the "3" rule column.

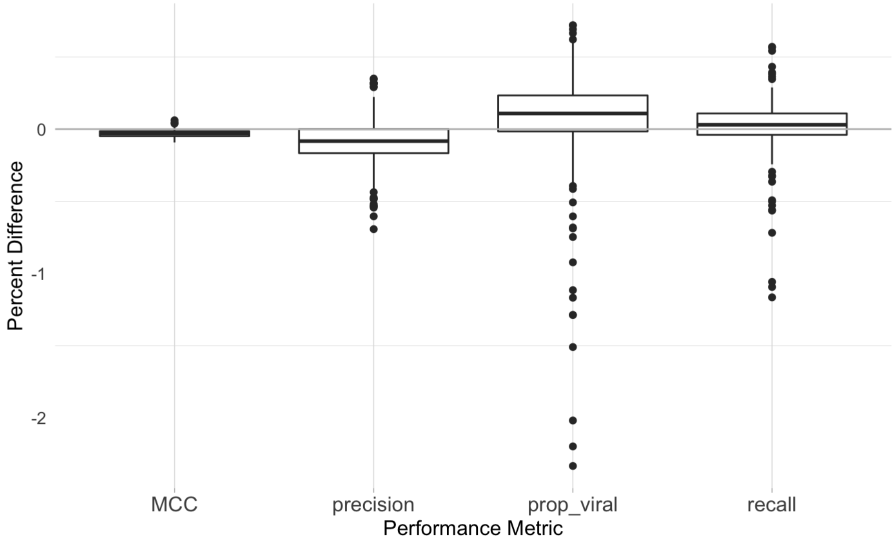


**Figure S4.** Difference between the performance metrics for the rulesets with and without the +0.5 subrules of the four single-tool rules. A percent difference greater than 0 means that the performance metric was bigger with the +0.5 subrules.

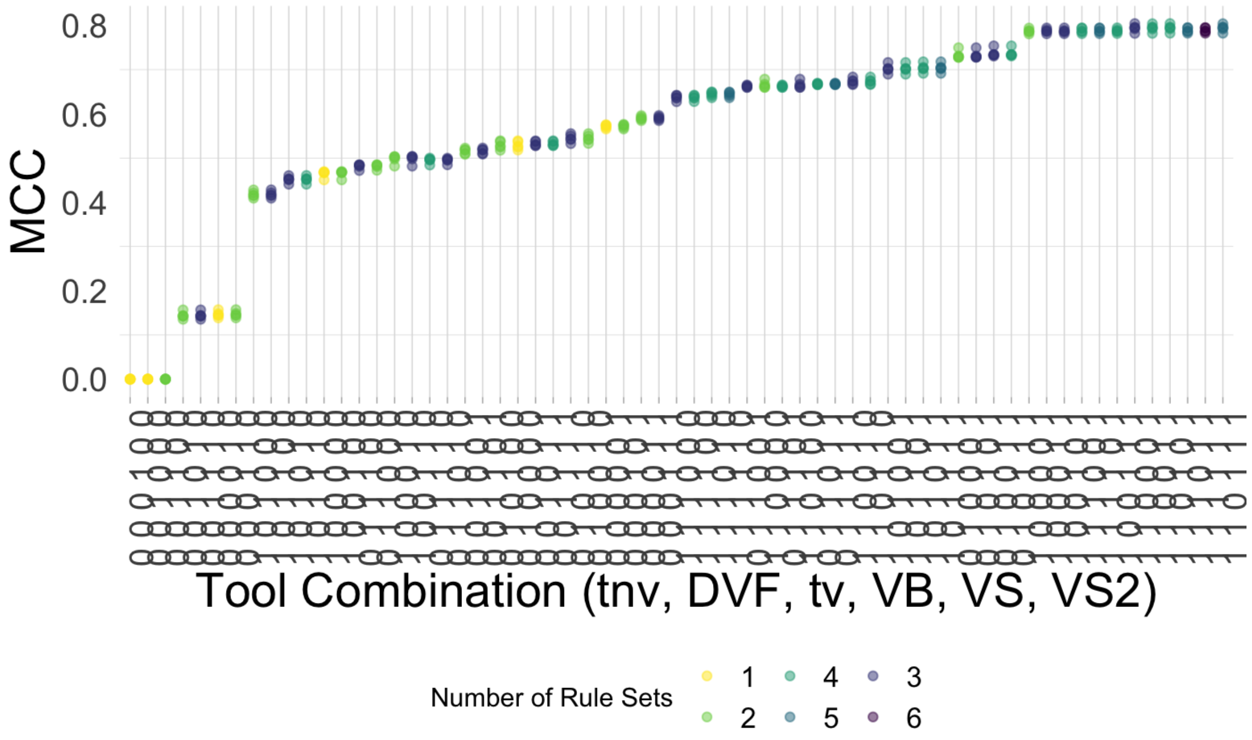


**Figure S5**. MCC for the 3-5 kb subset. Points are colored by the number of viral identification tools used in the ruleset and ordered by increasing MCC.

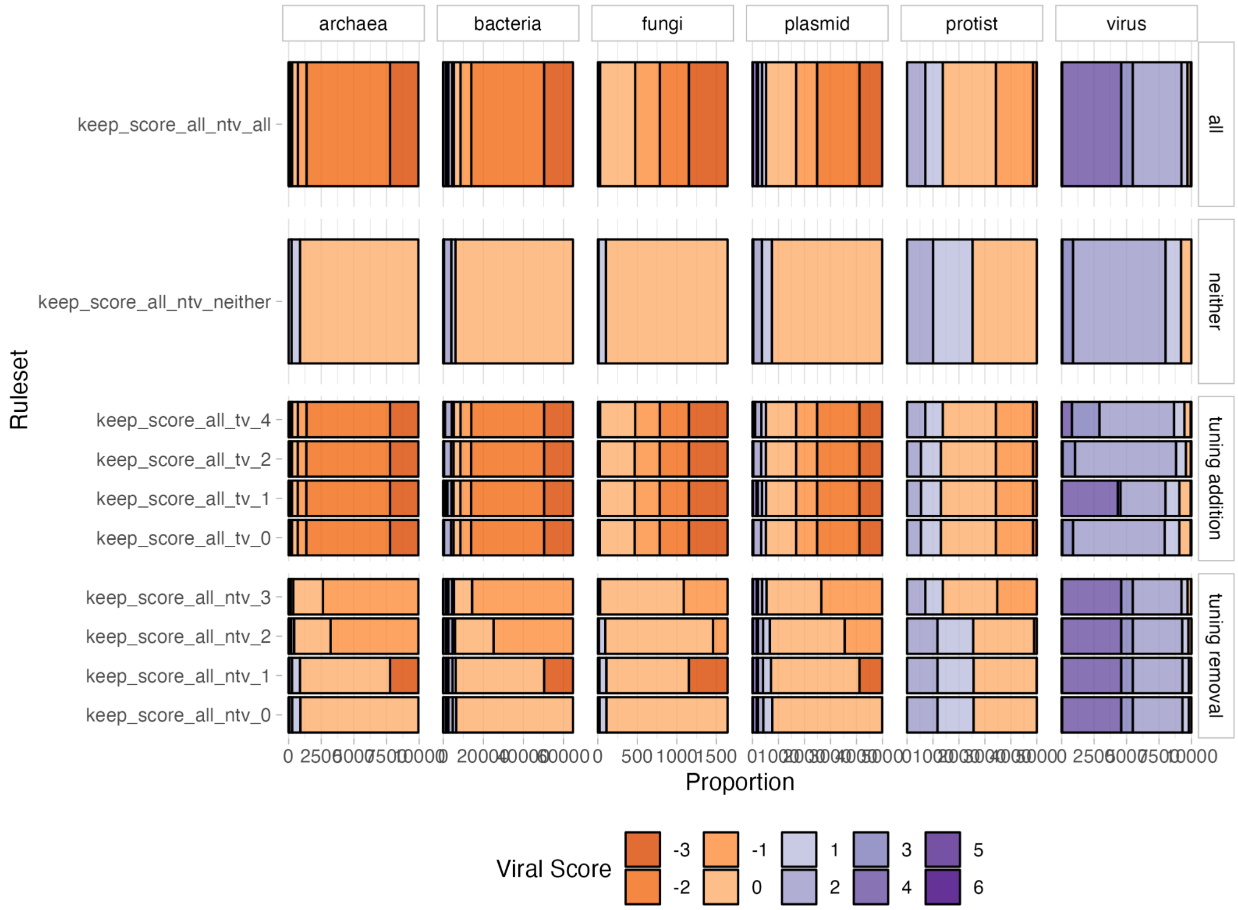


**Figure S6. Individual performance of each tuning rule.** Number of sequences with a given viral score for the **(Top)** tuning addition and **(Bottom)** tuning removal rules faceted by the sequence type. The tuning addition rules are (1) VirSorter2 viral hallmark genes > 2, (2) called viral by Kaiju and Kaiju match ration ≥ 0.3, (3) percent unknown ≤ 75% and sequence length ≤ 50 kb, and (4) percent viral ≥ 50% by VirSorter2 or CheckV. The tuning removal rules are (1) CheckV host genes > 50 and not called a provirus, (2) CheckV viral genes = 0 and host genes ≥ 1, (3) 3 times the number of CheckV viral genes*3 ≤ the number of CheckV host genes and not a provirus, (4) longer than 500 kb and one or fewer VirSorter2 hallmark viral genes, and (5) DeepVirFinder score ≤ 0.7 and p-value ≤ 0.05.

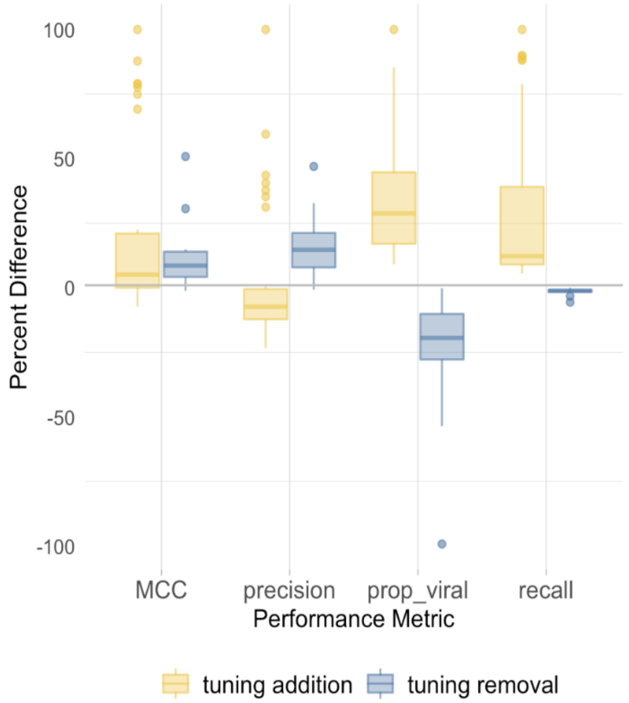


**Figure S7.** Overview of the change in MCC, precision, performance, and proportion viral on ruleset with the inclusion of the tuning removal and tuning addition rules. The percent difference is greater than zero if the performance metric is higher with the tuning rules.

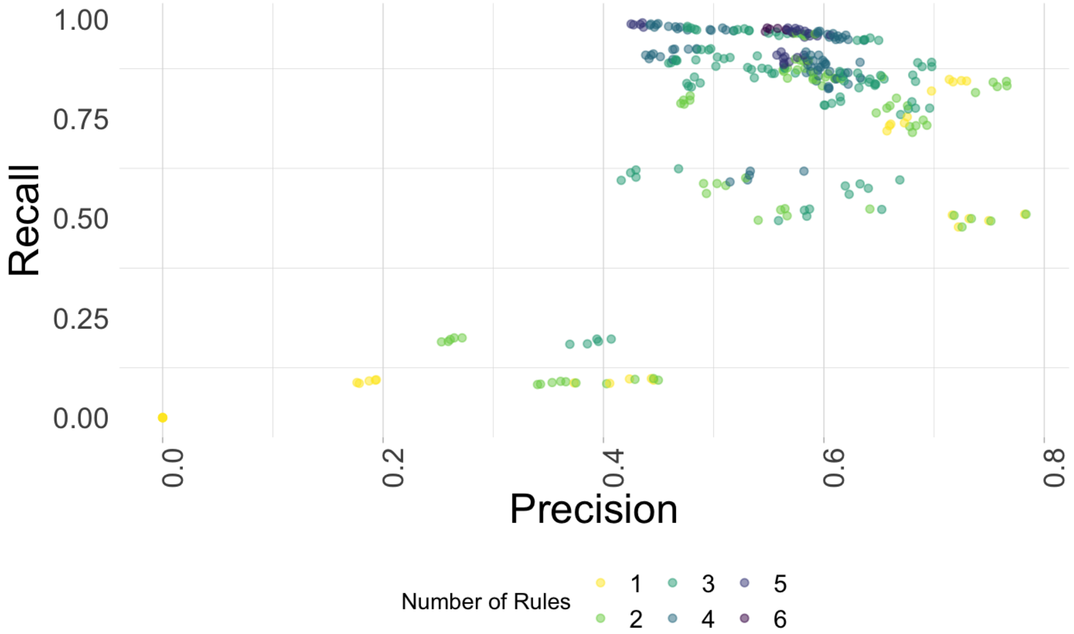


**Figure S8.** Scatter plot of recall and precision of virus versus non-virus on mock environmental metagenomic datasets. Each point represents a combination of tools run on a single testing set. Points are colored by how many rules were used.

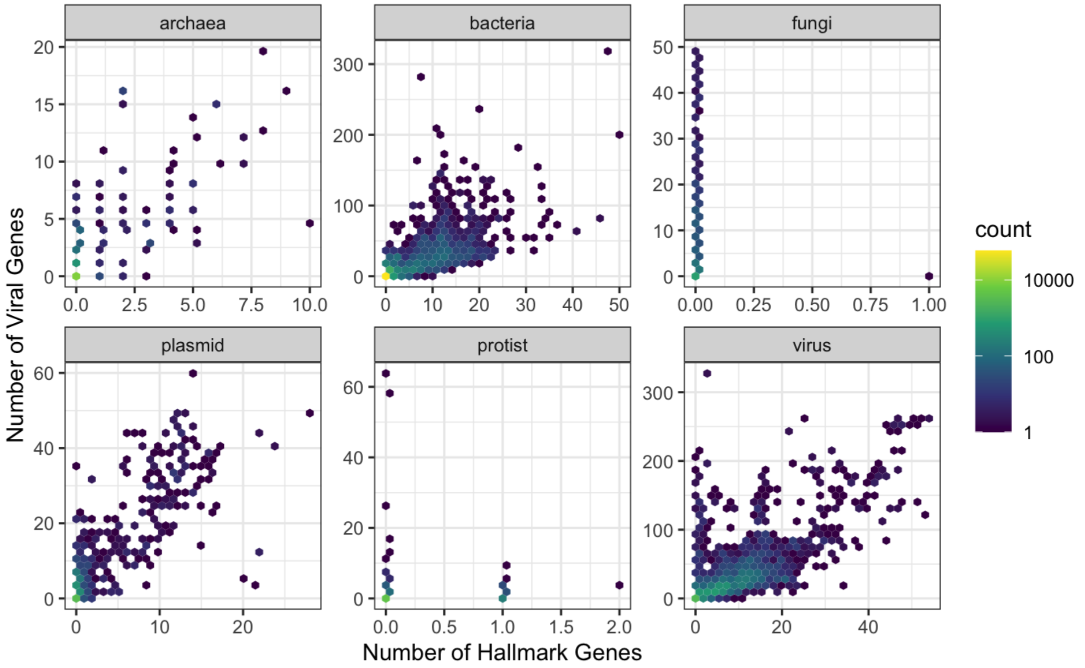


**Figure S9**. Comparison of number of hallmark genes (VirSorter2) and number of viral genes (CheckV) faceted by sequence type.

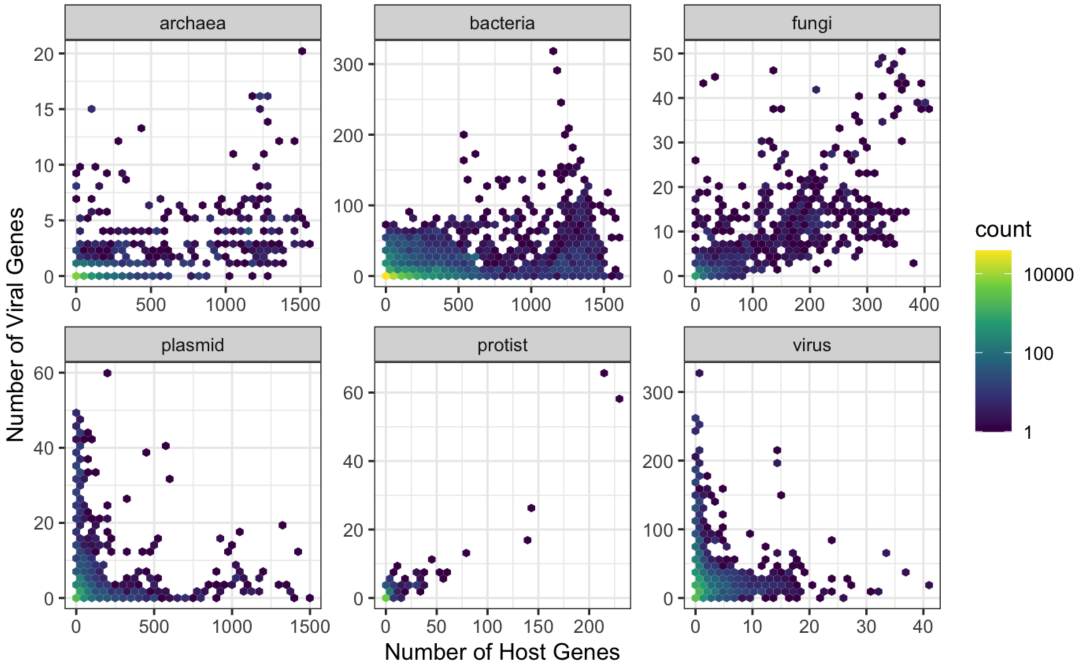


**Figure S10**. Comparison of number of host genes (CheckV) and number of viral genes (CheckV) faceted by sequence type.

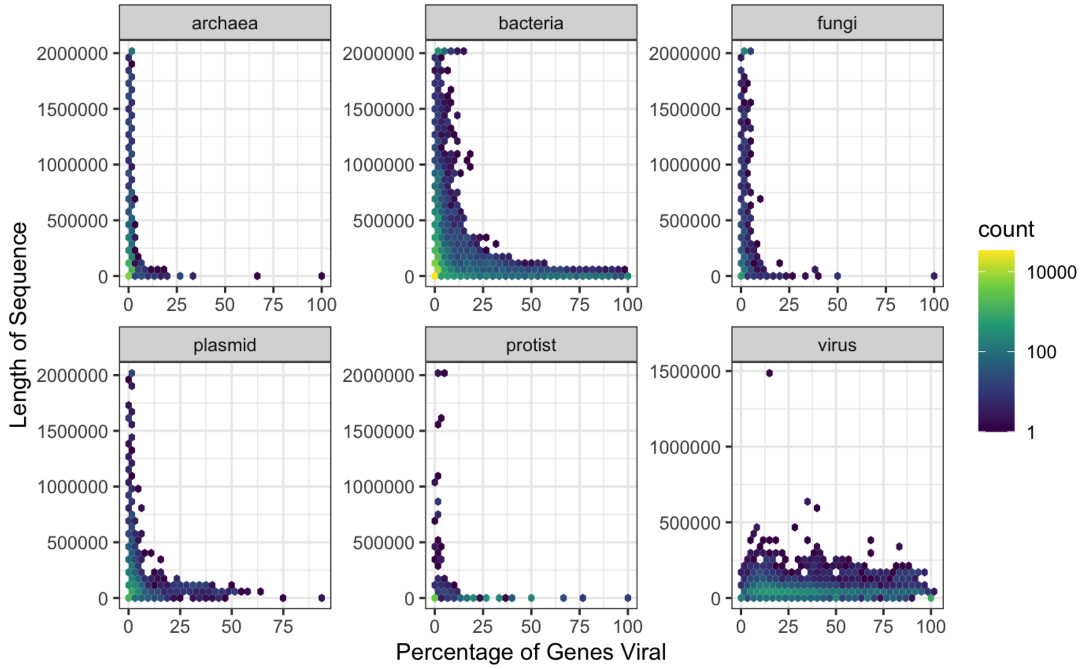


**Figure S11**. Comparison of percentage of genes viral (CheckV) and length of sequence faceted by sequence type.

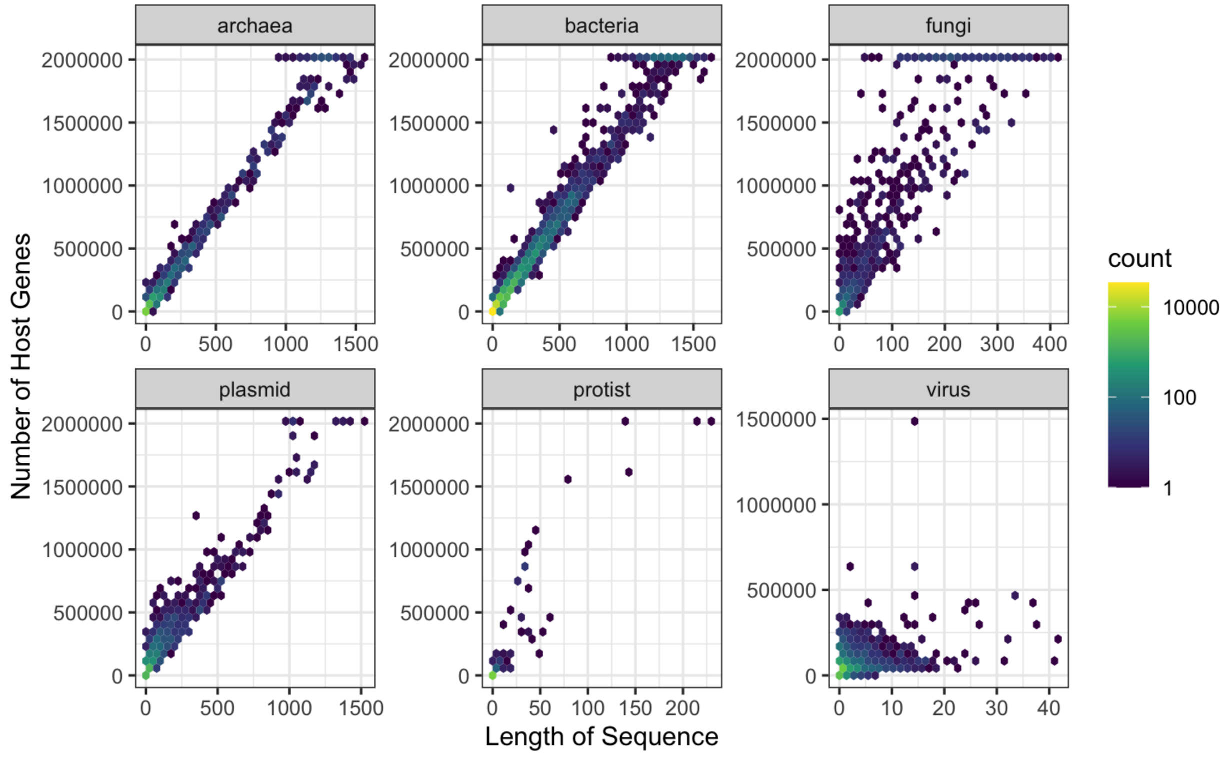


**Figure S12**. Comparison of length of sequence and number of host genes (CheckV) faceted by sequence type.

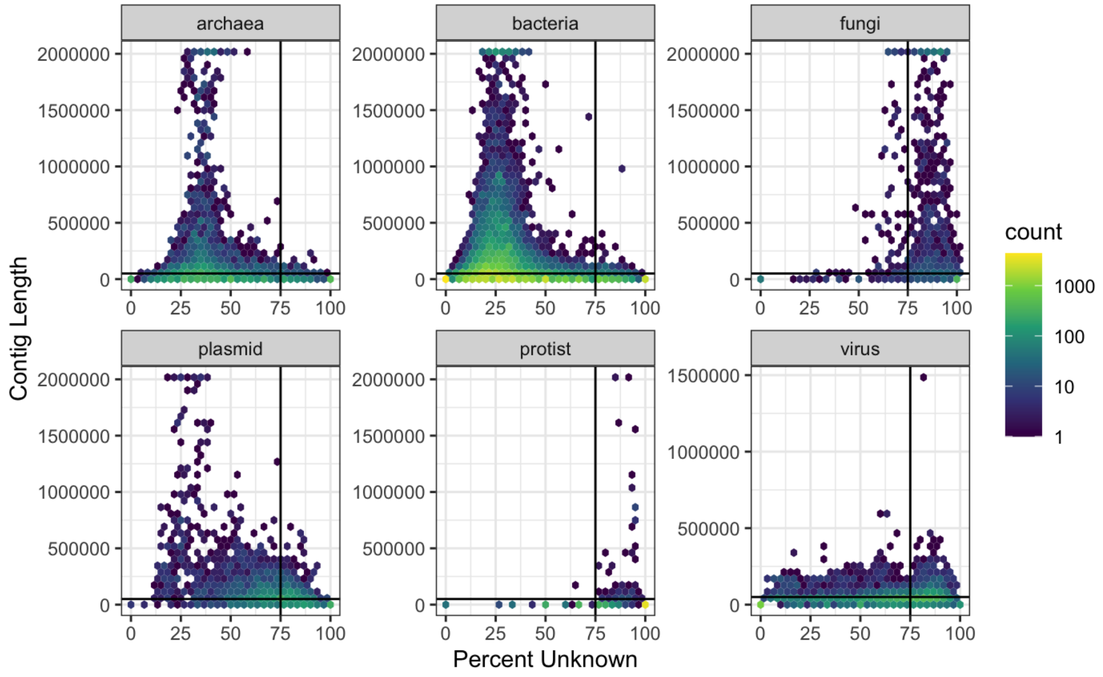


**Figure S13**. Comparison of percent unknown (CheckV) and contig length faceted by sequence type.

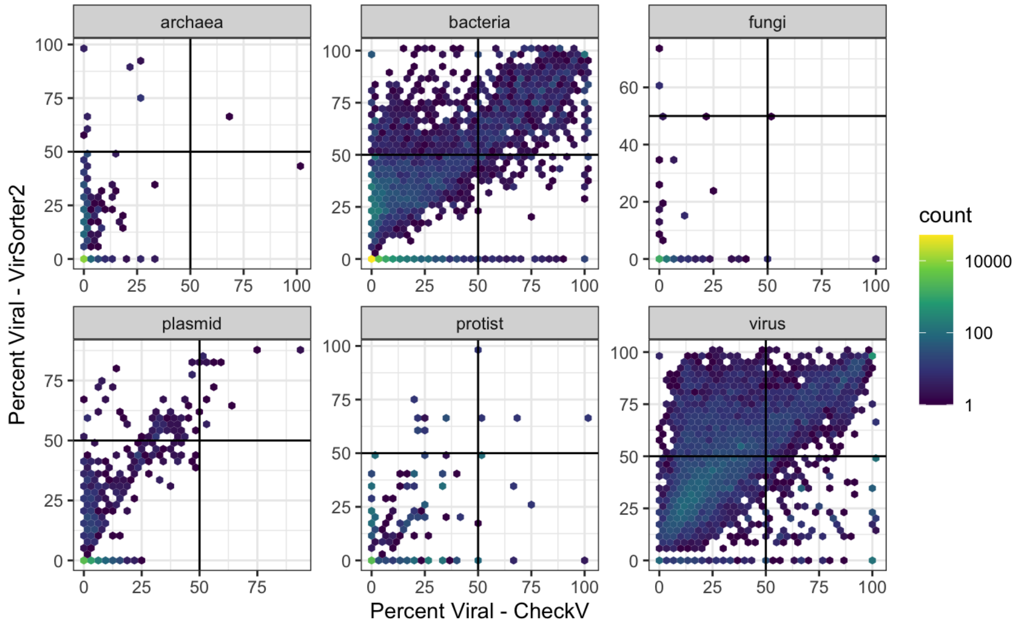


**Figure S14**. Comparison of percent viral (CheckV) and percent viral (VirSorter2) faceted by sequence type.

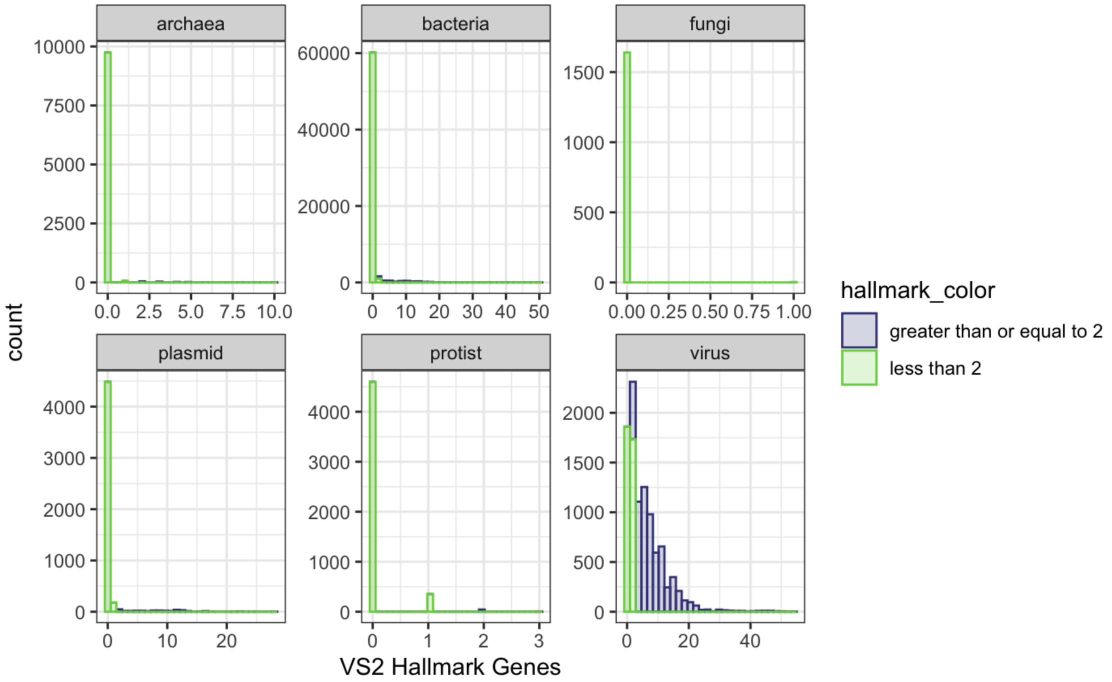


**Figure S15**. Histogram of number of hallmark genes (VirSorter2) faceted by sequence type and split by having more or less than 2 hallmark genes.

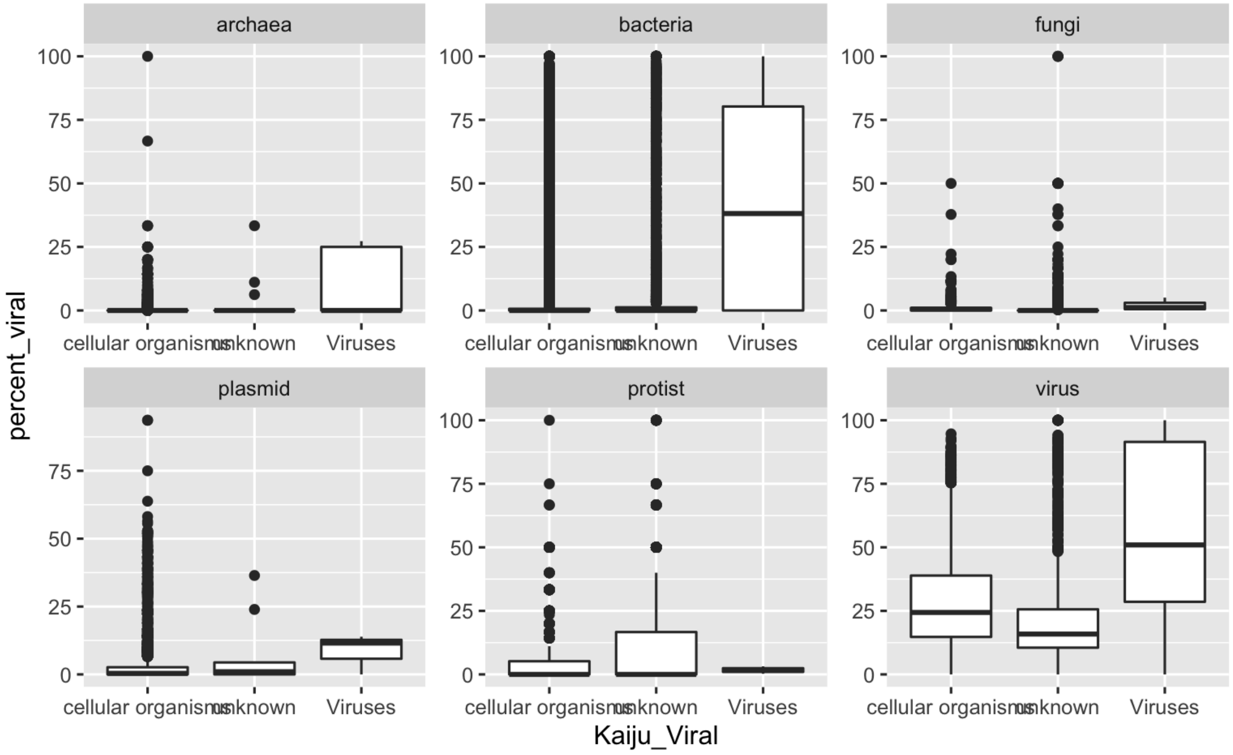


**Figure S16**. Comparison of Kaiju taxonomic assignment and percent viral (CheckV). All plots are faceted by true sequence type.

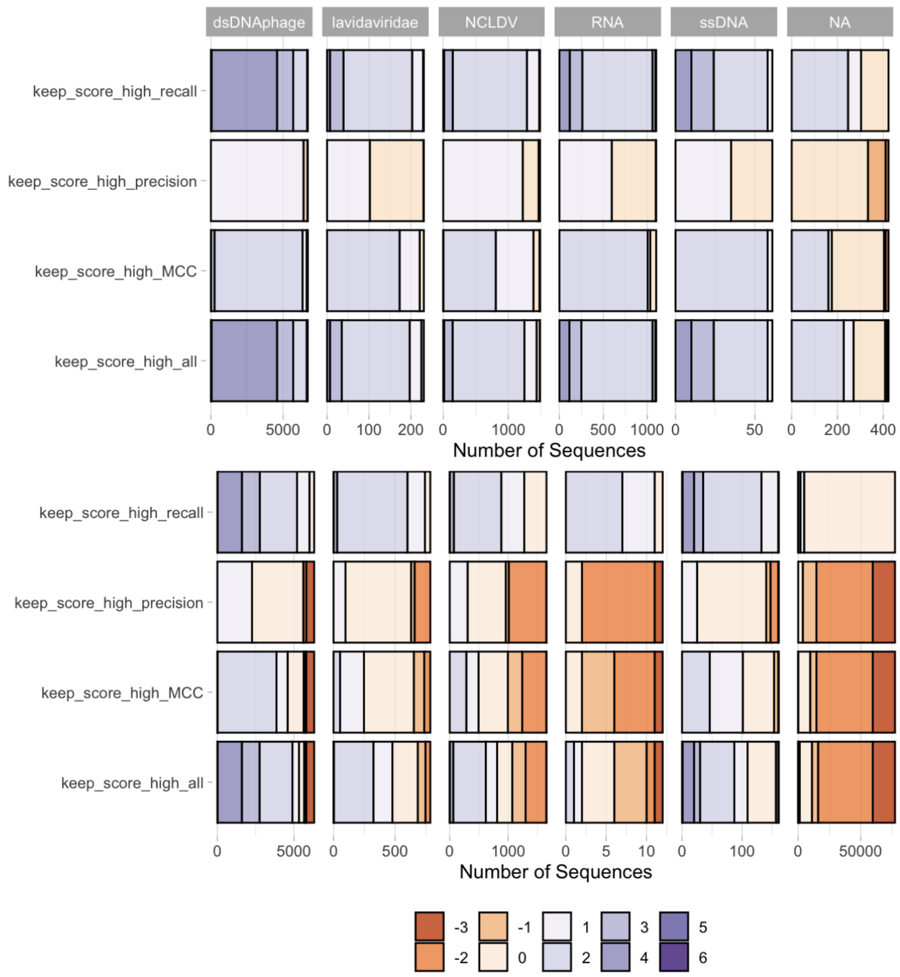


**Figure S17. Performance of four representative rulesets across viral types.** Number of sequences with a given viral score for four representative rulesets (high recall, high precision, high MCC, and all) faceted by the VirSorter2 viral group (NA represents sequences not called viral by VirSorter2) and separated based on **(A)** virus and **(B)** non-virus sequences. Note that the x-axes vary between panels and facets.

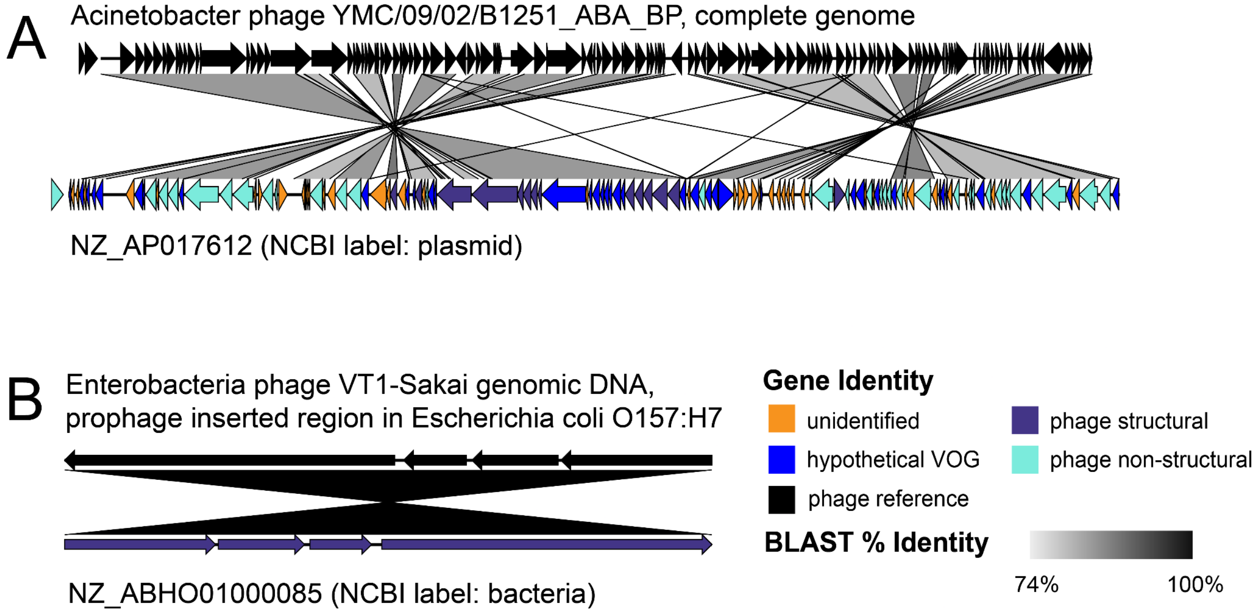


**Figure S18** - Sequence synteny plots indicating sequence similarity between three representative "false positive" sequences from the testing set (i.e., taxonomically assigned Bacteria in NCBI, but identified as viral by our highest MCC rulesetoptimized process) and their top NCBI Blast hit. (A) NZ_LLFE01000196 (NCBI label: bacteria) versus "Acinetobacter phage YMC/09/02/B1251_ABA_BP, complete genome". (B) NZ_AP017612 (NCBI label: plasmid) versus Salmonella phage SSU5. (C) NZ_ABHO01000085 (NCBI label: bacteria) versus "Enterobacteria phage VT1-Sakai genomic DNA, prophage inserted region in *Escherichia coli* O157:H7". All genes of the testing set sequences are labeled by their gene identity (orange: unidentified, blue: hypothetical VOG, purple: phage structural, and bright blue: phage non-structural. Greyscale bar represent BLAST percent identity score.

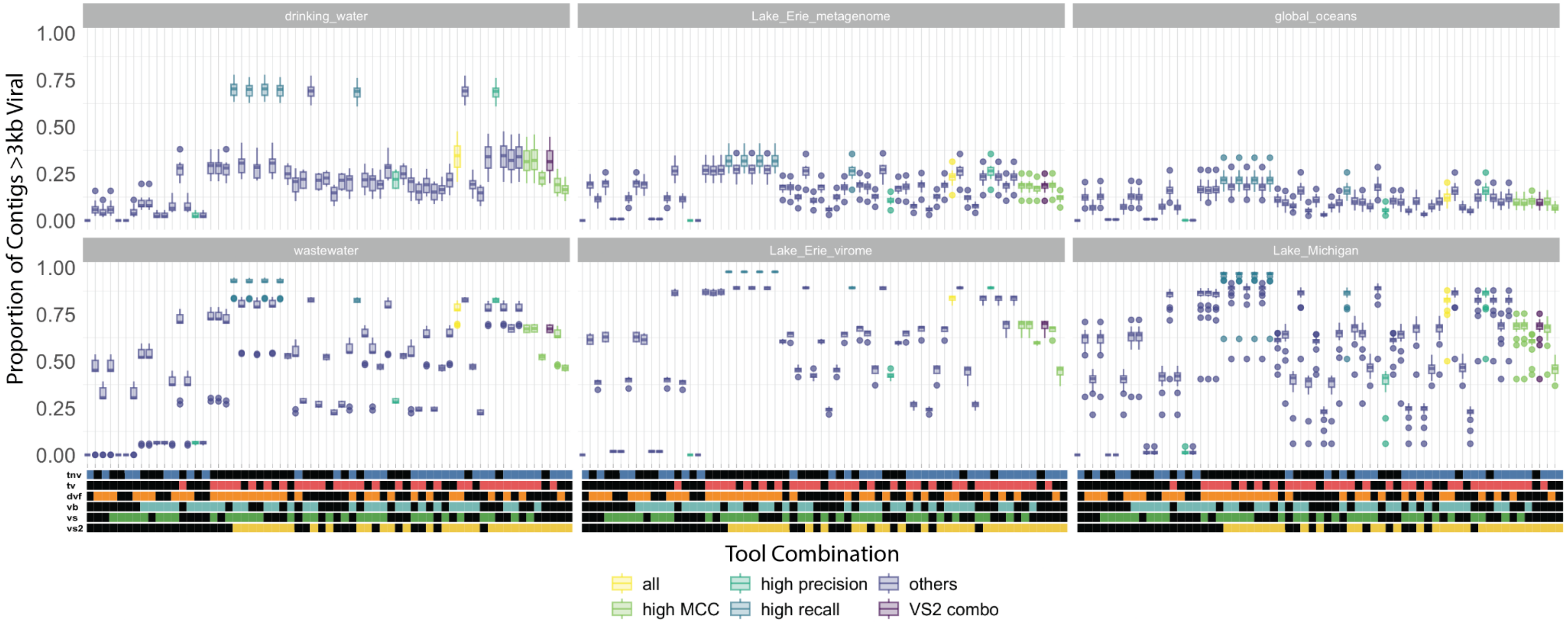
drinking_water
Lake_Erie_metagenome
global_oceans
wastewater
Lake_Erie_virome
Lake_Michigan
Proportion of Contigs >3kb Viral
1.00
0.75
0.50
0.25
0.00
tnv
tv
dvf
vb
vs
vs2
Tool Combination
all
high MCC
high precision
high recall
others
VS2 combo

**Figure S19**. **Proportion of viruses predicted by each tool combination across five environmental datasets.** Select tool combinations are colored to highlight their accuracy on the testing set. Multiple sets had the same MCC/precision/recall accounting for multiple t-tests with Bonferroni, and are colored the same accordingly. Vs2+tnv and vs2 were in both the high MCC and high precision groups. Tool combinations are sorted left-to-right by increasing MCC as shown in Figure X.
